# Induction of a Müller glial-specific protective pathway safeguards the retina from diabetes induced damage

**DOI:** 10.1101/2024.06.10.598362

**Authors:** Cheng-Hui Lin, Man-Ru Wu, Bogdan Tanasa, Praveen Prakhar, Alexander E. Davis, Liang Li, Alexander Xia, Yang Shan, Patrice E. Fort, Sui Wang

**Author notes:** **Please send correspondence to**: Sui Wang 1651 Page Mill Rd. Rm2114 Palo Alto, CA, 94304, USA (01)-650-586-9135. The authors have declared that no conflict of interest exists.

## Abstract

Diabetes can lead to cell-type-specific responses in the retina, including vascular lesions, glial dysfunction and neurodegeneration, all of which contribute to retinopathy. However, the molecular mechanisms underlying these cell type-specific responses, and the cell types that are sensitive to diabetes have not been fully elucidated. Employing single cell transcriptomic analyses, we profiled the transcriptional changes induced by diabetes in different retinal cell types in diabetic rat models as the disease progressed. Rod photoreceptors, a subtype of amacrine interneurons, and Müller glial cells exhibited rapid responses to diabetes at the transcript levels. Genes associated with ion regulation were upregulated in all three cell types, suggesting a common response to diabetes. Furthermore, focused studies revealed that while Müller glial cells initially increased the expression of genes playing protective roles, they cannot sustain this beneficial effect as the disease progressed. We explored one of the candidate protective genes, Zinc finger protein 36 homolog (Zfp36), and observed that depleting *Zfp36* in rat Müller glial cells in vivo using AAV-based tools exacerbated early diabetes-induced phenotypes, including gliosis, neurodegeneration, and vascular defects. Notably, the over-expression of *Zfp36* slowed the development of phenotypes associated with diabetic retinopathy. In summary, this work unveiled retinal cell types that are sensitive to diabetes and demonstrated that Müller glial cells can mount protective responses through *Zfp36*. The failure to maintain *Zfp36* levels contributes to the development of diabetic retinopathy.

## Introduction

Diabetic retinopathy (DR) affects over 7 million individuals in the US, a number anticipated to rise with the increasing prevalence of diabetes^1^. The imperative to better understand this disease and devise effective therapeutic approaches is pressing. Despite notable successes, existing treatment modalities, such as anti-VEGF drugs, steroids, vitreous opacities removal, and photocoagulation, possess limitations and side effects. The need for repetitive intraocular injections of anti-VEGF drugs or steroids, coupled with the non-responsiveness of a considerable patient subset^2^, poses challenges. Moreover, the potential harm to retinal neurons from prolonged anti-VEGF drug use, given VEGF’s recognized neuronal protective effects, is a concern^3,4^. Photocoagulation and surgeries primarily offer palliative relief. All these interventions target the advanced stages of DR. They can slow down the progress of DR but fail to restore vision loss. One of the promising ways to safeguard vision under diabetic conditions is by intercepting the initiation or impeding the early progression of DR before vision is compromised. Yet, the molecular mechanisms governing the onset and early progression of DR remain poorly understood.

The vertebrate retina comprises diverse neurons, glial cells, and vascular components. Diabetes triggers cell-type-specific responses in the retina at early stages, evident in both animal models and non-proliferative diabetic retinopathy (NPDR) patients^5–7^. Specifically, diabetic-induced degeneration and atrophy of retinal ganglion cells (RGCs) were observed in postmortem retinas and various rodent models at early stages^5,8–12^. Diabetes also affects photoreceptors, evidenced by impaired color vision and reduced contrast sensitivity at early stages^13,14^. Photoreceptors can further contribute to DR development through mechanisms like releasing proinflammatory proteins^15–17^. Retinal macroglial cells - Müller glia (MG) and astrocytes – as well as microglia swiftly respond to diabetes, undergoing morphological and functional changes^6^. While their roles in the development of DR are acknowledged, precise functions remain to be fully elucidated. The retinal vasculature responds to diabetes with lesions such as microaneurysms, hemorrhages, and capillary defects - early NPDR signs in humans^18^. In rodent models, endothelial dysfunction and pericyte loss were observed and possibly underly the diabetes-induced vascular defects^19^. These findings demonstrate that diabetes can elicit diverse responses in different cell types, and these cell types probably interact with each other and collectively contribute to the development of DR. In this study, we sought to unveil the molecular mechanisms underlying the diabetes-induced cell-type-specific responses, with the hope of advancing our understanding of DR and informing therapeutic strategies.

To elucidate cell-type-specific molecular mechanisms, an optimal strategy involves labeling and isolating individual retinal cell types, followed by profiling their molecular alterations at various levels. Unfortunately, the lack of molecular tools for in vivo labeling and manipulation of diverse retinal cell types has significantly impeded such investigations. Recent strides in single-cell transcriptomic analyses offer a promising solution to this challenge^20^. Single-cell RNA-seq facilitates straightforward and robust exploration of the transcriptomes of thousands or even millions of individual cells within a tissue. Leveraging gene expression profiles, cells can be effectively clustered into distinct types, obviating the need for labeling or isolating specific cell types. This approach allows for a comprehensive examination of transcriptional changes associated with diabetes-induced responses in a multiplexed manner.

In this study, we conducted single-cell RNA-seq (scRNA-seq) analyses on rat retinas at 1, 2, or 3 months following the induction of diabetes. Among the 53 identified retinal cell types, we particularly observed noteworthy transcriptomic alterations in rod photoreceptors, a specific subtype of amacrine cells, and Müller glial cells. Notably, the upregulated genes in these cell types were found to be primarily associated with ion regulation. Subsequent focused investigations revealed that Müller glial cells play a crucial role in exerting protective and beneficial effects during the early stages of DR. The failure of this adaptive response may contribute to the initiation and progression of DR.

## Results

### scRNA-seq analyses reveal diabetes-induced cell-type-specific retinal responses in the early stages of DR

The retinal responses induced by diabetes have been studied using a variety of diabetic rodent models that show phenotypes reminiscent of human DR at early stages^21^. Among these models, the streptozotocin (STZ)-induced diabetic rat model has been well characterized and widely used to study the effect of diabetes on the retina. STZ is a chemical that can selectively depletes insulin-secreting beta cells in the pancreas and has long been used to mimic type 1 diabetes in rodents^22^. When injected with a single low dose of STZ (65mg/g body weight), more than 90% of male rats developed high fasting blood glucose levels (>250mg/dL) within 2 days (Figure S1A). Male rats with fasting glucose levels consistently higher than 250mg/dL were considered diabetic. Ninety percent of these diabetic male rats remained diabetic for at least 5 months, which were used in this study. Insulin was not used to avoid confounding factors.

We profiled diabetes-induced gene expression changes in the rat retina at different stages after STZ injection via scRNA-seq. The retinas were harvested, rapidly dissociated and submitted for scRNA-seq (10x Genomics) at 1, 2 or 3 months after STZ injection (Figure 1A and S1A). Biological replicates were included for both control and STZ retinas at each stage, and total of 12 samples were sequenced. Approximately 7335 to 13759 cells were successfully captured and sequenced for each condition (Figure S1B). Based on well-established markers for retinal cell types^23^, we assigned known cell identities to the 53 clusters grouped by an unbiased clustering method (Seurat 4.1) (Figure 1B-C & S1C-F). The top 24 clusters with more than 500 cells were selected for downstream analyses to ensure robust statistical analyses (Figure S2A-B). These included all the major retinal neuronal types (rod, cone, horizontal, bipolar, amacrine and ganglion cells) and Müller glial cells (Figure 1C & S1C-D). However, these did not include subtypes of retinal ganglion cells (RGCs), which were not observed in sufficient numbers in our study and would have required a specific enrichment approach. Retinal astrocytes, microglia and endothelial cells were not included in follow-up in-depth analyses due to the low number of cells captured.

**Figure 1.**
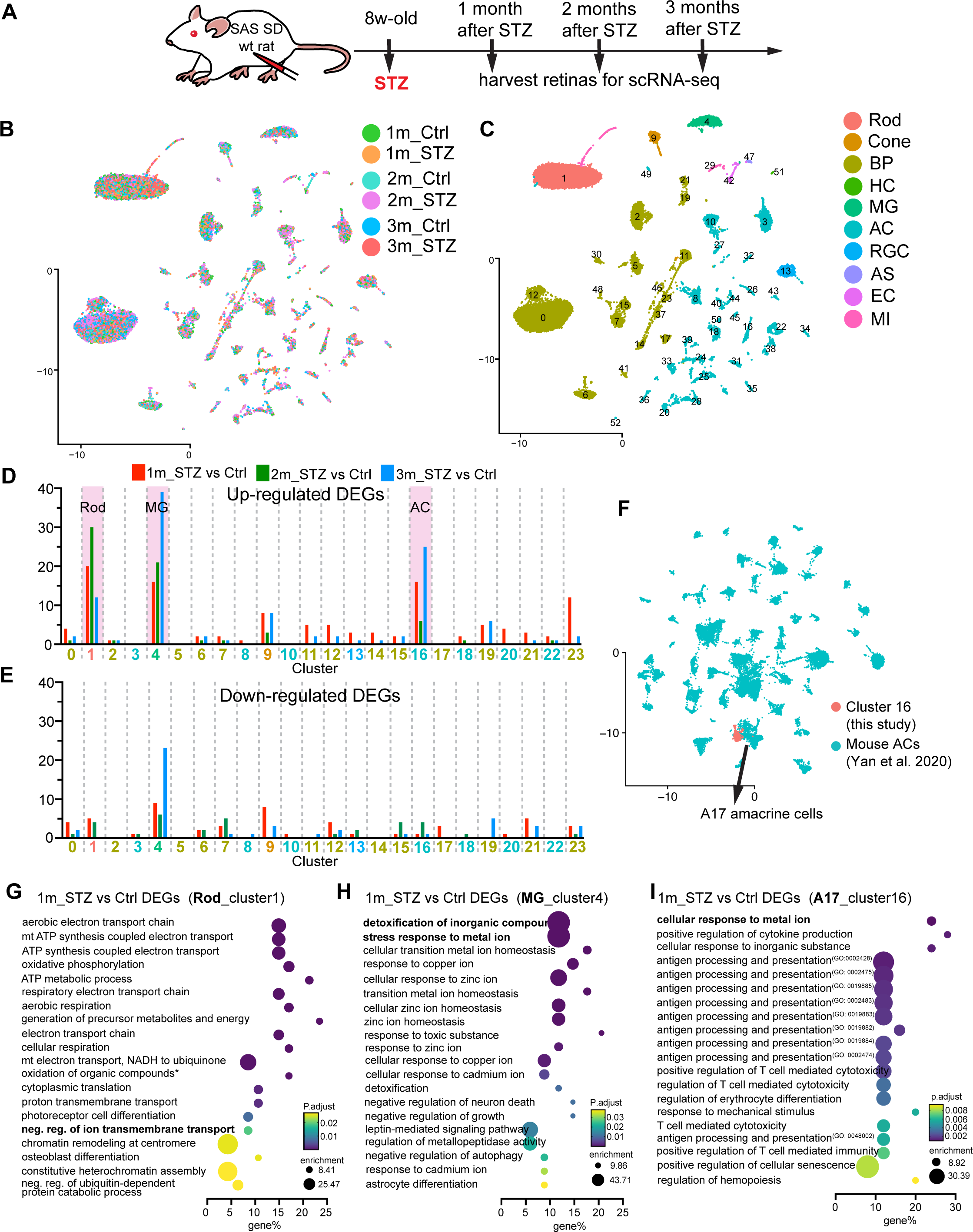
scRNA-seq reveals retinal cell types are sensitive to diabetes at the transcript level. **(A)** Schematic representation of the experimental design. **(B-C)** UMAP plots illustrating scRNA-seq data. Conditions are highlighted in (B), and cell types are indicated in (C). BP: bipolar cells; HC: horizontal cells; MG: Müller glial cells; AC: amacrine cells; RGC: retinal ganglion cells; AS: astrocyte; EC: endothelial cells; MI: microglia. **(D-E)** Number of upregulated (D) or downregulated (E) DEGs (|log2(Fold Change)|>1, P<0.05) per cluster. X-axes represent cluster numbers, with cluster types indicated by colors as shown in (C): Red for 1m_STZ vs 1m_Ctrl, Green for 2m_STZ vs 2m_Ctrl, and Blue for 3m_STZ vs 3m_Ctrl. **(F)** Query of cluster 16 on published reference datasets of amacrine cells (Yan et al., 2020). **(G-I)** GO enrichment analyses of DEGs in MG, Rod, and A17 cells between 1m_STZ and 1m_Ctrl conditions. The top 20 GO terms are displayed.

We first investigated which cell clusters exhibited significant changes at the transcript level in response to diabetes by quantifying the number of differentially expressed genes (DEGs) between STZ and control retinas (Figure 1D-E). At 1, 2, and 3 months after STZ injection, rod photoreceptor (cluster 1), Müller glial cells (cluster 4), and a subtype of amacrine cells (cluster 16) consistently showed higher counts of up-regulated DEGs in STZ retinas compared to the other clusters. This suggests that these 3 retinal cell types are sensitive to diabetes at the transcript level in the early stages of DR. Interestingly, we observed a limited number of down-regulated DEGs in most cell clusters (Figure 1E). Furthermore, to determine the identity of cluster 16 amacrine cells, we performed a comparison between cluster 16 cells and the various subtypes of mouse amacrine cells previously identified by scRNA-seq^24^ (Figure 1F). The gene expression profiles of cluster 16 cells matched well with those of A17 mouse amacrine cells, which are known to be a key component of the scotopic vision (hereafter, designated as A17 amacrine cells)^25^.

We further performed gene ontology (GO) enrichment analysis for the DEGs in rods, A17 amacrine cells and Müller glia to uncover possible underlying mechanisms. At 1 month post STZ injection, rod photoreceptors primarily upregulated genes involved in energy metabolism and oxidative phosphorylation (Figure 1G). A17 amacrine cells and Müller glia mainly upregulated genes involved in ion regulation (Figure 1H-I). Interestingly, regulation of ion transport is also among the top 20 enriched GO terms in rod photoreceptors (Figure 1G). This suggests that cellular responses to ions could be common early responses induced by diabetes in retinal cells.

As the disease progresses, rods, A17 amacrine cells and Müller glia upregulated genes involved in different pathways. Rods continued upregulating genes involved in energy related pathways at 2 months post STZ, such as purine ribonucleotide biosynthetic process (Figure S2C). However, this effect was not observed at 3 months after STZ injection since enriched GO terms were not found by analyzing the upregulated DEGs in rods. While we didn’t detect any enrich GO terms in A17 amacrine cells at 2 months post STZ, they significantly upregulated genes involved in regulating responses to biotic stimulus and defense responses at 3 months post STZ (Figure S2D). Müller glial cells started to upregulate genes involved in the regulation of vascular associated smooth muscle cell proliferation at 2 months post STZ, and genes involved in blood coagulation and angiogenesis at 3 months post STZ (Figure S2E-F).

Overall, scRNA-seq analyses revealed that rods, A17 amacrine cells and Müller glia are sensitive to diabetes at the transcript level at early stages. One of the common early responses induced by diabetes at the transcript level could be the upregulation of genes involved in ion regulation. Apart from this common response, rod, A17 amacrine cells, and Müller glia regulated genes involved in different biological pathways in response to diabetes.

### Retinal Müller glial cells upregulate potential protective genes in response to diabetes but failed to sustain this beneficial effect

Müller glial cells are known to respond to insults of diverse etiologies in the retina^26^. They have been shown to be one of the first responders to diabetes in the retina^27^ and showed most significant changes at the transcript level in our scRNA-seq analyses (Figure 1D). We therefore conducted comprehensive studies to explore the diabetes-induced responses in Müller glial cells and to uncover the molecular mechanisms driving these responses.

We examined DEGs in the Müller glial cell cluster (cluster 4) at 1, 2, or 3 months post-STZ injection in our scRNA-seq dataset (Figure 2A). Notably, most DEGs upregulated at 1 month did not sustain upregulation with disease progression, referred to as “up-down DEGs”, as shown in the heatmap (Figure 2B). Approximately 72% of these up-down DEGs exhibited enriched expression specifically in Müller glial cells (Figure 2B). We scrutinized the published literature for potential gene functions of each DEG, most of which had not been previously studied in the retina. Based on their roles in glial cells and other systems, we categorized them into four groups (Figure S3). "Protective" genes encompass those with established roles in anti-inflammatory, neuroprotective, pro-survival, anti-apoptotic, anti-proliferative, anti-oxidative stress, or anti-angiogenic pathways. "Negligent" genes are those involved in pro-inflammatory, pro-apoptotic, pro-migratory, or oncogenic pathways. "Context-dependent" genes can exhibit protective or negligent roles depending on the cellular context. Genes with unclear roles were labeled as "unclear”. Interestingly, 22 out of 36 up-down DEGs were potential protective genes (Figure 2B and Figure S3), suggesting Müller glial cells primarily upregulate protective pathways in response to diabetes at early stages.

**Figure 2.**
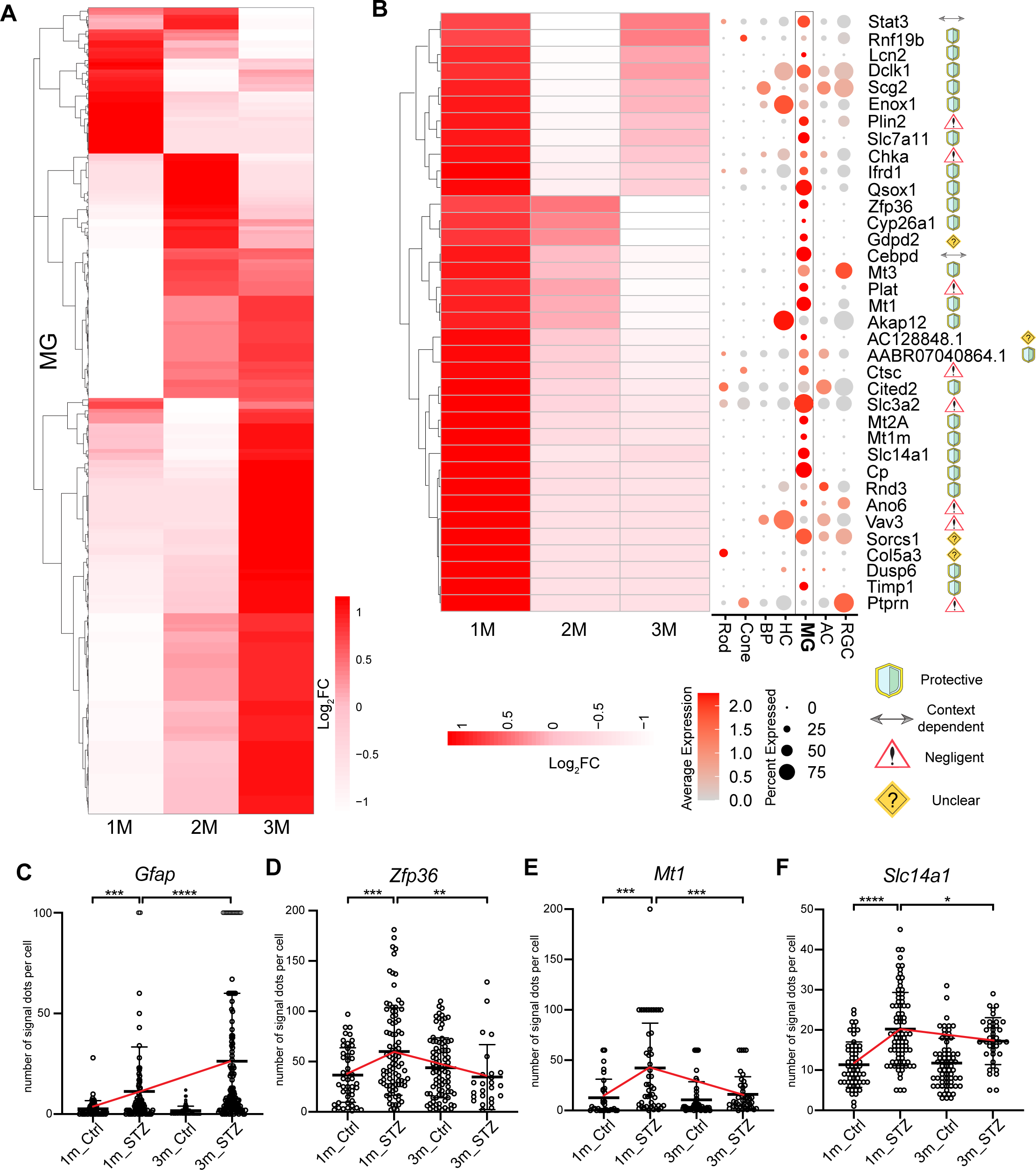
Müller glial cells upregulate protective genes but fail to maintain the beneficial effects as DR progresses. **(A)** Heatmap illustrating the log2(fold changes) of DEGs in Müller glial cells (MG) at 1, 2, or 3 months after citrate buffer or STZ injection (P<0.2). Each row represents a gene. **(B)** Analysis of genes temporarily upregulated at 1 month (1m_STZ vs 1m_Ctrl). Left: Heatmap presenting log2(fold changes) of DEGs; Middle: Dotplot showcasing expression in major retinal cell types; Right: Gene names and predicted roles. Legends are provided at the bottom. **(C-F)** Quantification of mRNA levels of the positive candidate gene *Gfap* (C) and candidate protective genes *Zfp36*, *Mt1*, and *Slc14a1* (D-F) in Müller glial cells (MG), as revealed by smFISH. Each dot represents a single MG cell (identified by Sox9 antibody or in situ signals; also refer to Figure S4) freshly dissociated from rat retinas. The Y-axis denotes the number of mRNA signal dots in each cell. Mean ± SD. Unpaired t-test with Welch’s correction (two-tailed) was applied for statistical analysis. Significance levels are denoted as follows: * P < 0.05, ** P < 0.01, *** P < 0.001, **** P < 0.0001. N >= 3 rats.

We then validated the expression of some of the candidate protective genes through single molecule fluorescent RNA in situ hybridization (smFISH), which enables quantitative assessment of mRNA levels in a cell-type-specific manner. The number of smFISH signal dots exhibits a strong correlation with mRNA levels^28,29^. The *Gfap* gene, which was continuously upregulated in Müller glial cells as the disease progresses in STZ retinas, served as a positive control ^27^(Figure 2C and S4A-C). Three candidate protective gene *Zfp36*, *Mt1*, and *Slc14a1* were validated. These genes were selected for validation due to their high enrichment in Müller glial cells, as indicated by our scRNA-seq data (Figure 2B). Furthermore, they serve as representatives of genes crucial in anti-inflammation, anti-oxidative stress, or neuroprotective pathways, respectively. To enhance precise quantification, we swiftly dissociated rat retinas into single cell preparations at 1 or 3 months post citrate buffer or STZ injection and conducted smFISH targeting *Gfap*, *Zfp36*, *Mt1*, or *Slc14a1*. Sox9, a well-known Müller glia marker, was co-detected using either antibodies or smFISH probe targeting *Sox9* mRNAs to label Müller glial cells. We quantified the number of smFISH signal dots (correlating with mRNA levels) for *Gfap*, *Zfp36, Mt1* or *Slc14a1* in both Sox9+ Müller glial cells and Sox9-retinal cells. The smFISH signal dot counts for *Gfap* mRNA significantly increased in Müller glial cells at 1 month after STZ injection compared to citrate buffer-injected control retinas. This elevation was further enhanced at 3 months post STZ injection, aligning with the sustained upregulation of *Gfap* mRNA levels observed in our scRNA-seq data (Figure S4A-C). In contrast, the number of smFISH signal dots for *Zfp36*, *Mt1*, or *Slc14a1* mRNA in Müller glial cells showed an initial increase at 1 month post STZ injection compared to citrate buffer-injected controls. However, this increase diminished at 3 months post STZ injection (Figure 3D-F, 4A, S4). In addition, the smFISH signal dot counts for all three genes in Sox9-cells showed no significant changes between STZ and control retinas, suggesting a specific regulation of these genes in Müller glial cells in response to diabetes (Figure S4C-E). We hence concluded that Müller glial cells transiently upregulated the candidate protective genes *Zfp36*, *Mt1*, and *Slc14a1* at 1 month post STZ, yet failed to sustain this regulation by the 3-month post STZ.

**Figure 3.**
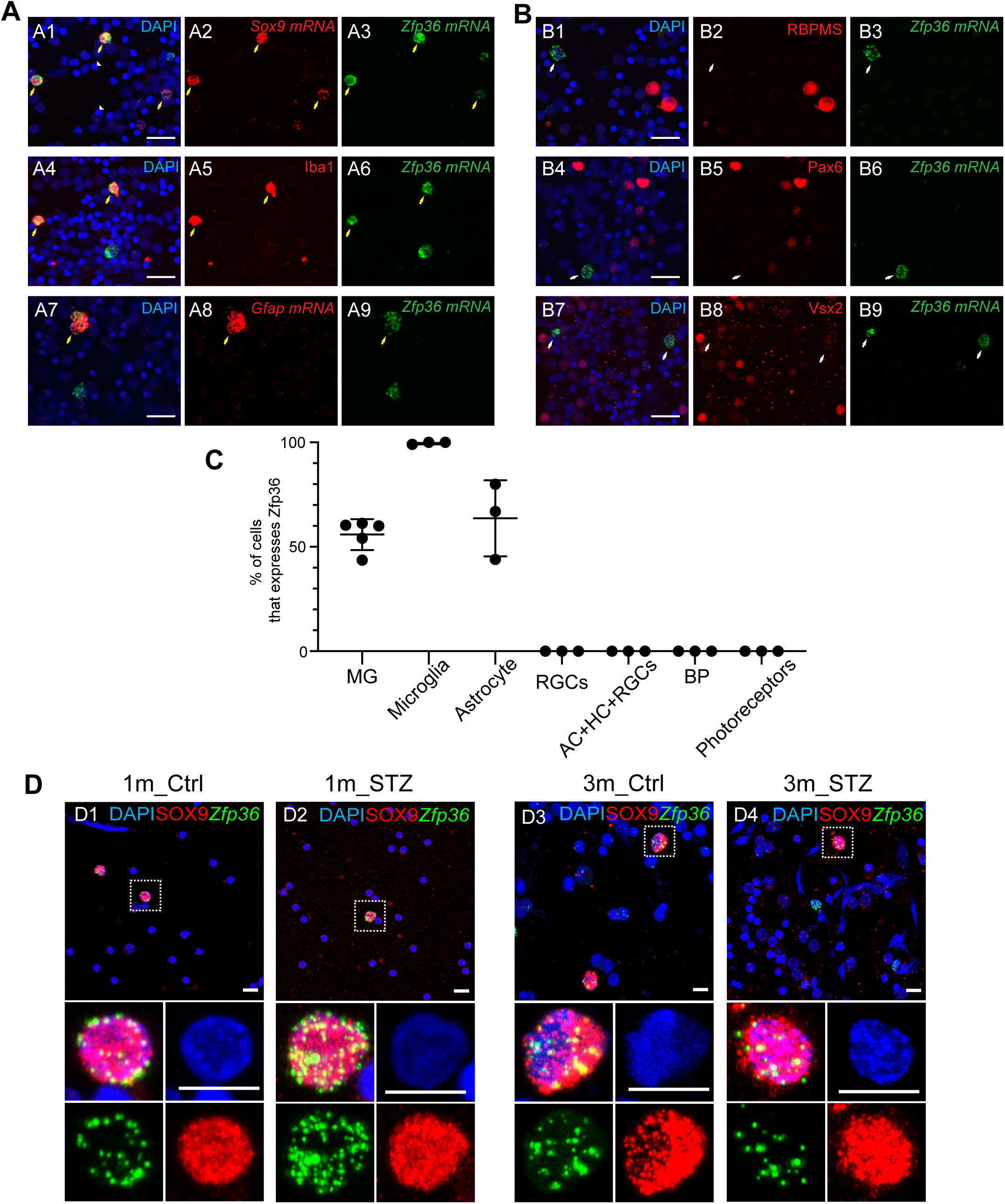
Zfp36 is expressed in retinal glial cells, and transiently upregulated in Müller glial cells. **(A-B)** Co-detection of Zfp36 mRNA by FISH along with established markers for specific retinal cell types. *Sox9* mRNA: predominantly in MG (A1-A3). Iba1 antibody staining: microglia (A4-A6). Gfap mRNA: astrocytes (A7-A9). RBPMS antibody staining: retinal ganglion cells (RGCs, B1-B3). Pax6 antibody staining: amacrine cells, horizontal cells, and RGCs (B4-B6). Vsx2 (Chx10) antibody staining: bipolar cells (B7-B9). Scale bar: 20 μm. Yellow arrows indicate cells co-expressing *Zfp36* and cell-type-specific markers, while white arrows indicate cells expressing *Zfp36* but not the indicated markers. Triangles (A1) represent photoreceptors. **(C)** Percentage of indicated cell types expressing *Zfp36*. **(D)** Temporal upregulation of *Zfp36* in Sox9+ rat MG at 1 month after STZ injection. Red: Sox9 antibody staining. Green: *Zfp36* mRNA in situ. Scale bar: 10 μm.

We further investigated whether the gene expression changes induced in Müller glial cells by diabetes were comparable to those triggered by other forms of insults. Hoang et al. previously documented gene expression changes at the single-cell level in Müller glial cells using an N-methyl-D-aspartate (NMDA)-induced acute retinal injury model^30^.

We juxtaposed the gene expression profile of Müller glial cells in our diabetic rat model with that of Müller glial cells in the published NMDA-induced acute retinal injury mouse model^30^ (Figure S5A). The transcript profiles of Müller glial cells in diabetic rats exhibited overall similarity with those of mouse Müller glial cells at 24 to 72 hours post NMDA injection, as shown in the UMAP plot (Figure S5B-D). Notably, 19 of the 36 up-down DEGs outlined in Figure 2B exhibited a parallel temporal upregulation in the NMDA injury model during the early stages (3-12 hours post NMDA injection), followed by subsequent downregulation at 24-72 hours post NMDA injection, including genes like *Mt1* and *Slc14a1* (Figure S5E). This suggests that Müller glial cells may engage common pathways in response to both diabetes and acute injury.

In summary, through analysis and validation of scRNA-seq data, we showed that Müller glial cells initially upregulate genes with potential protective effects; however, they fail to sustain this beneficial effect as the disease advances. Additionally, our investigation unveiled potential genes that may participate in common pathways in response to various insults.

### Depletion of *Zfp36* in Müller glial cells accelerates the development of DR

We performed targeted studies to further investigate the potential protective roles of Müller glial cells in response to diabetes at early stages by elucidating the roles of candidate up-down DEGs. Notably, we focused on the *Zfp36* (*Zinc finger protein 36 homolog*) gene, recognized for its regulatory functions in various systems. Encoding an RNA-binding protein, Zfp36 binds to AU-rich elements in the 3’UTR of numerous pro-inflammatory mRNAs, leading to swift mRNA degradation^31^. While acknowledged as a potent inhibitor of inflammation-related pathways in macrophages^31^, its role in the retina remains unclear. Given the key role of inflammation in the development of DR^32^, we investigated whether Zfp36 expressed in MG exerts protective effects in the retina in response to diabetes.

We first verified the expression pattern of *Zfp36* in the retina. At the transcript level, Zfp36 mRNAs were identified in approximately 55% of Müller glial cells, 100% of microglial cells, and 60% of astrocytes in wildtype rat retinas as detected by smFISH (Figure 3A-C). No smFISH signal dots above background were detected in retinal photoreceptor, horizontal, bipolar, amacrine, and retinal ganglion cells (Figure 3B-C). Additionally, although *Zfp36* mRNA levels per Müller glial cell exhibited temporary upregulation at 1 month post-STZ, there was no apparent increase in the percentage of Müller glial cells expressing Zfp36. At the protein level, the commercially available Zfp36 antibody was ineffective for immunohistochemistry on rat retinal cryo-sections in our experiments. Furthermore, obtaining a sufficient amount of Müller glial cells for western blotting analyses via fluorescence-activated cell sorting (FACS) proved challenging. Consequently, we investigated Zfp36 protein alterations using lysates from the entire rat retina. Consistent with scRNA-seq and smFISH data, Zfp36 protein levels exhibited upregulation at 1 month post STZ injection and subsequent downregulation at 3 months post-STZ injection compared to controls injected with citrate buffer (Figure S6A). Notably, while it remains unknown if the early Müller glial-response also takes place in humans, the transient upregulation of *Zfp36* observed in STZ-treated rat retinas aligns with the trends of *Zfp36* regulation in human donor samples. Like what was seen in 1-month STZ rat samples, *Zfp36* levels were increased in tissues from diabetic donors without DR when compared to matched non-diabetic donors. Like what was seen in 3-month STZ rat samples, *Zfp36* levels were decreased in tissues from diabetic donors with NPDR when compared to matched non-diabetic donors (Figure S6B). Overall, in response to diabetes, Zfp36 mRNA and protein levels exhibited a transient upregulation in Müller glial cells. However, this adaptive response was not sustained as DR advanced.

To elucidate the role of Zfp36 in DR, we performed in vivo loss-of-function analyses in the rat retina using Adeno-Associated Virus (AAV)-based tools capable of selectively driving expression in Müller glial cells. We previously have demonstrated that the AAV7m8-pGFAP-EGFP (AAV7m8 capsid) viruses can direct gene expression specifically in mouse Müller glial cells in vivo through intravitreal injection, achieving approximately 95% of specificity and 55% efficiency^33^. We first evaluated the performance of this AAV in rats in vivo. Following intravitreal injection into postnatal day 4 (P4) rat pups, the AAV successfully labeled approximately 60% of Müller glial cells with a 95% specificity in adult rats, and the sustained expression of EGFP persisted for at least 5 months (Figure S7). Subsequently, we designed 4 microRNA-based shRNA cassettes to target rat *Zfp36* mRNA. One of them (Zfp36shRNA-1) could reduce rat *Zfp36* mRNA levels by at least 80% (Figure S8A-B). This cassette was integrated into the 3’UTR of the *Egfp* gene within the AAV-pGFAP-EGFP backbone. The AAV7m8 capsid was then used to package the resulting AAV7m8-pGFAP-EGFP-Zfp36shRNA-1 viruses (AAV7m8-Zfp36KD) for subsequent knock-down experiments. Control experiments employed AAVs expressing a control shRNA targeting the bacterial *lacZ* gene (which does not exist in mammalian cells) (AAV7m8-Ctrl) (Figure 4A).

**Figure 4.**
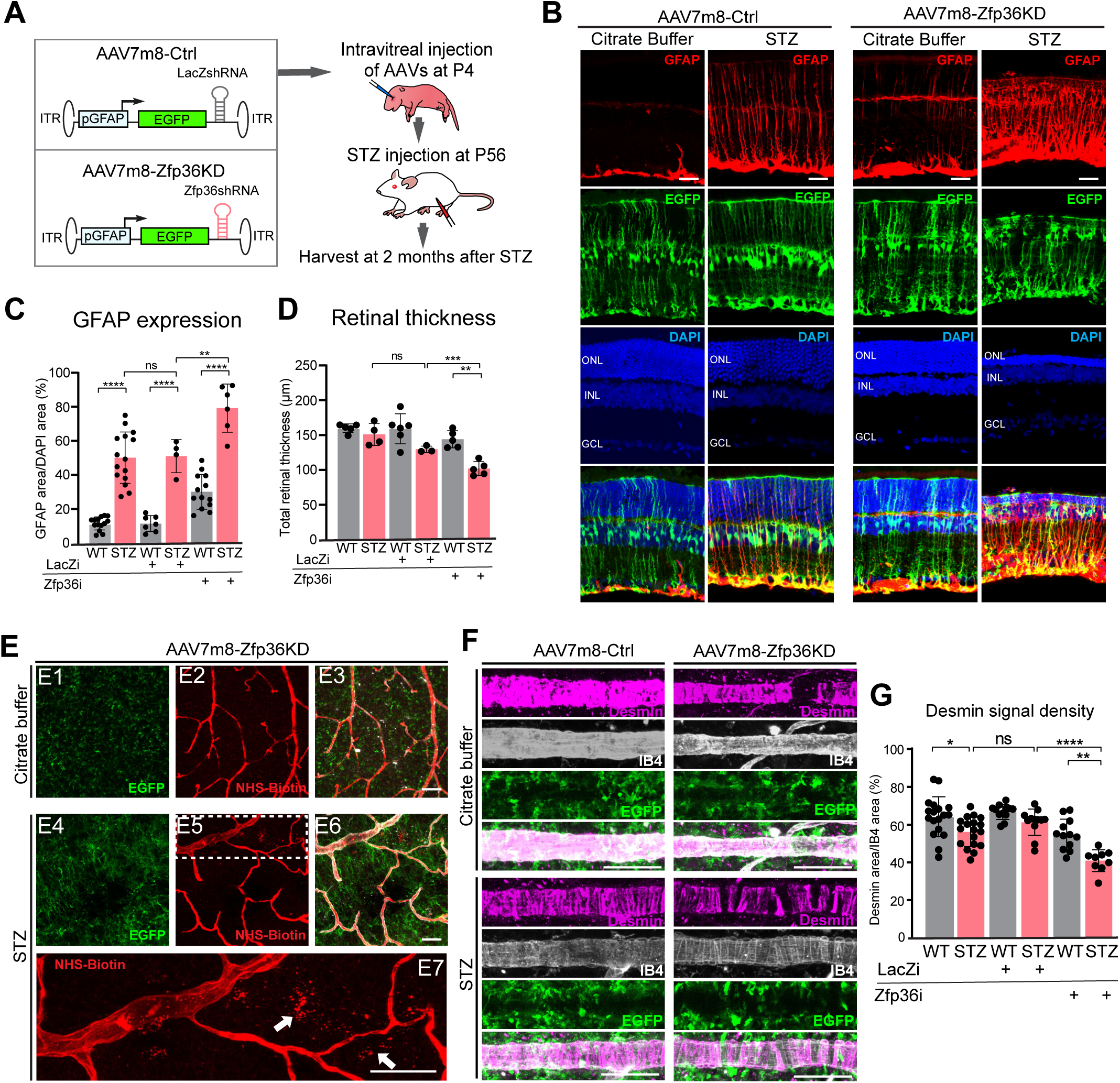
Depletion of *Zfp36* in Müller glial cells accelerates the development of DR. **(A)** Experimental design schematic. **(B)** Retinal sections following administration of AAV7m8-Ctrl or AAV7m8-Zfp36KD viruses. Red: GFAP antibody staining. Scale bar: 20 μm. **(C-D)** Quantification of GFAP expression (percentage of GFAP+ area over DAPI+ area) and total retinal thickness in control and STZ retinas. Mean ± SD. Unpaired t-test with Welch’s correction (two-tailed). ** P < 0.01, *** P < 0.001, **** P < 0.0001, ns: P >= 0.05. N numbers indicated in the charts. LacZi: rats received AAV7m8-Ctrl. Zfp36i: rats received AAV7m8-Zfp36KD. **(E)** Retina vessel permeability assay in rats receiving AAV7m8-Zfp36KD. E1-E3: rats also received citrate buffer. E4-E7: rats also received STZ. Red: NHS-biotin tracer detected by Alexa Fluor 555 conjugated-Streptavidin. Green: EGFP signals from AAVs. E7 is a higher magnification of the highlighted region in E5. White arrows indicate NHS-biotin leakage from vessels in STZ retinas. Scale bar: 20 μm. **(F)** Representative images of Desmin antibody staining. **(G)** Quantification of desmin signal density. LacZi: rats received AAV7m8-Ctrl. Zfp36i: rats received AAV7m8-Zfp36KD. Mean ± SD. Unpaired t-test with Welch’s correction (two-tailed). * P < 0.05, ** P < 0.01, *** P < 0.001, **** P < 0.0001, ns P >= 0.05. N numbers indicated in the charts.

We injected AAV7m8-Zfp36KD or AAV7m8-Ctrl into wild type rat retinas via the intravitreal route at P4. Choosing this early time point enhances transduction efficiency in rat retinas compared to injections in adults, likely attributed to the smaller globe size of the rat eye at P4. Subsequently, at 8 weeks of age, these rats received either STZ or citrate buffer, as illustrated in Figure 4A. Following a 2-month interval post-STZ/citrate buffer injection, we harvested and examined the retinas.

In the absence of diabetes, the injection of control AAV7m8-Ctrl along with citrate buffer did not elicit any noticeable gross changes in rat retinas in vivo (Figure 4B). Knocking down *Zfp36* in rat Müller glial cells via AAV7m8-Zfp36KD along with citrate buffer did not result in detectable developmental defects. However, it induced mild Müller glia gliosis in control rats, as evidenced by the upregulation of GFAP, a hallmark of gliosis. This observation suggests that Zfp36 plays a role in maintaining the quiescent state of rat Müller glial cells under wildtype conditions. In the presence of diabetes, rats that received AAV7m8-Zfp36KD exhibited significantly enhanced gliosis, evident through elevated GFAP expression, and increased neurodegeneration, as indicated by reduced retinal thickness, compared to diabetic rats received control AAV7m8-Ctrl (Figure 4B-D).

In addition, we investigated whether manipulating Zfp36 expression in Müller glial cells could lead to vascular defects, a crucial aspect of DR. In the absence of diabetes, neither AAV7m8-Ctrl nor AAV7m8-Zfp36-KD retinas displayed any retinal vascular leakage, as assessed by detection of the 226kDa fluorescently labeled NHS-biotin tracer^34^. However, upon depletion of *Zfp36* in Müller glial cells in diabetic rats, we observed tracer leakage from the vasculature in regions infected with AAV7m8-Zfp36KD (Figure 4E). Moreover, diabetic rats with AAV7m8-Zfp36KD injection exhibited a significant reduction in desmin (a well-established marker for vascular smooth muscle cells^35^) expression on the main retinal vessels compared to diabetic rats with AAV7m8-Ctrl injection, or non-diabetic controls (Figure 4F-G). These findings underscore the role of Müller glial-expressed Zfp36 in maintaining the proper function of Müller glial cells under normal conditions and safeguarding retinal neurons and vasculature under diabetic conditions.

To exclude potential off-target effects of Zfp36shRNA-1, we conducted rescue experiments by co-administering AAVs expressing a codon-optimized rat *Zfp36* resistant to Zfp36shRNA-1 (AAV7m8-ZR) along with AAV7m8-Zfp36KD in rats. The retinas of these rats exhibited a normal appearance (Figure S8C-D). Notably, no significant Müller glia gliosis, neurodegeneration, or vascular defects were observed in these retinas at 2 months post STZ injection.

Taken together, our findings demonstrate that targeted depletion of *Zfp36* in Müller glial cells accelerates the development of DR, suggesting that Müller glial cells can confer beneficial effects through the action of *Zfp36*.

### Enhanced Zfp36 expression in Müller glial cells protects the retina from damage induced by diabetes

We further explored the potential of elevating Zfp36 levels in Müller glial cells to mitigate DR progression in STZ-induced diabetic rats. To achieve this, we created an AAV co-expressing rat Zfp36 and EGFP in rat Müller glial cells in vivo (AAV7m8-Zfp36OE) (Figure 5A). AAV7m8-EGFP viruses, expressing EGFP alone, served as controls (Figure 5A).

**Figure 5.**
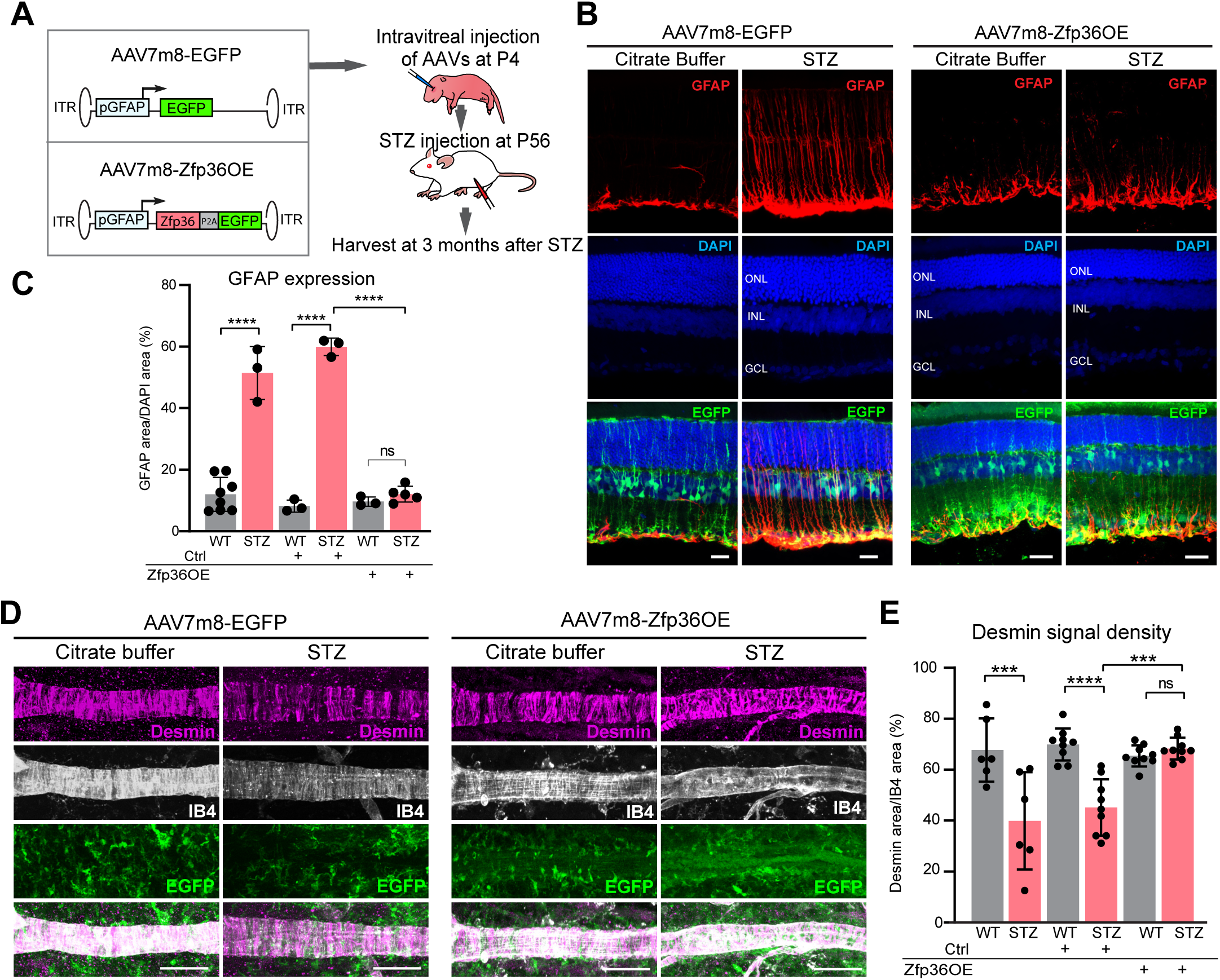
Sustained expression of *Zfp36* in Müller glial cells can protect the retina from diabetes-induced damage. **(A)** Experimental design schematic. **(B)** Retinal sections following administration of AAV7m8-EGFP or AAV7m8-Zfp36OE viruses. Red: GFAP antibody staining. Scale bar: 20 μm. **(C)** Quantification of GFAP expression (percentage of GFAP+ area over DAPI+ area) in control and diabetic retinas. Mean ± SD. Unpaired t-test with Welch’s correction (two-tailed). **** P < 0.0001, ns P >= 0.05. N numbers indicated in the charts. Ctrl: rats received AAV7m8-EGFP. Zfp36OE: rats received AAV7m8-Zfp36OE. **(D)** Representative images of desmin antibody staining. **(G)** Quantification of desmin signal density in the retina. Ctrl: rats received AAV7m8-EGFP. Zfp36OE: rats received AAV7m8-Zfp36OE. Mean ± SD. Unpaired t-test with Welch’s correction (two-tailed). *** P < 0.001, **** P < 0.0001, ns P >= 0.05. N numbers indicated in the charts.

At postnatal day 4, we intravitreally injected AAV7m8-Zfp36OE or AAV7m8-EGFP viruses to wild type rat pups, followed by either citrate buffer or STZ administration at 8 weeks of age. At the 3-month mark post-STZ injection, we harvested and examined these rat retinas. In the absence of diabetes, AAV7m8-Zfp36OE or AAV7m8-EGFP did not induce any retinal phenotypes (Figure 5B). Under diabetic conditions, the retinas of diabetic rats that received AAV7m8-EGFP exhibited Müller glia gliosis as expected, indicated by the upregulation of GFAP (Figure 5B). These retinas also showed reduced desmin expression in retinal main vessels compared to non-diabetic controls. Conversely, the retinas of diabetic rats that received AAV7m8-Zfp36OE showed a significant reduction of gliosis, and comparable desmin levels with non-diabetic controls (Figure 5B-E). The overall morphology of these retinas also appeared normal. Therefore, elevating Zfp36 expression in Müller glial cells emerges as a protective mechanism, shielding the retina from diabetes-induced damage.

## Discussion

We conducted a comprehensive examination of molecular responses in the retina induced by diabetes using scRNA-seq at three stages post diabetes induction in rats. Retinal rod photoreceptor cells, Müller glial cells, and A17 amacrine cells are highly sensitive to diabetes at the transcript level, displaying differentially expressed genes as early as 1 month post diabetes onset. In-depth analyses focused on Müller glial cells unveiled a previously unrecognized dynamic regulation of genes throughout the disease progression. Protective genes showed transient upregulation at early stages, but this beneficial effect waned as the disease advanced. Based on these findings, we propose a working model (Figure 6): in the presence of diabetes, Müller glial cells initiate the upregulation of protective genes early on to shield the retina from diabetes-induced damage. However, the adaptive response cannot be sustained as the disease progresses. To support this model, we demonstrated that augmenting the levels of Zfp36, one of the genes transiently upregulated in Müller glial cells, can effectively protect the retina from diabetes-induced damage.

**Figure 6.**
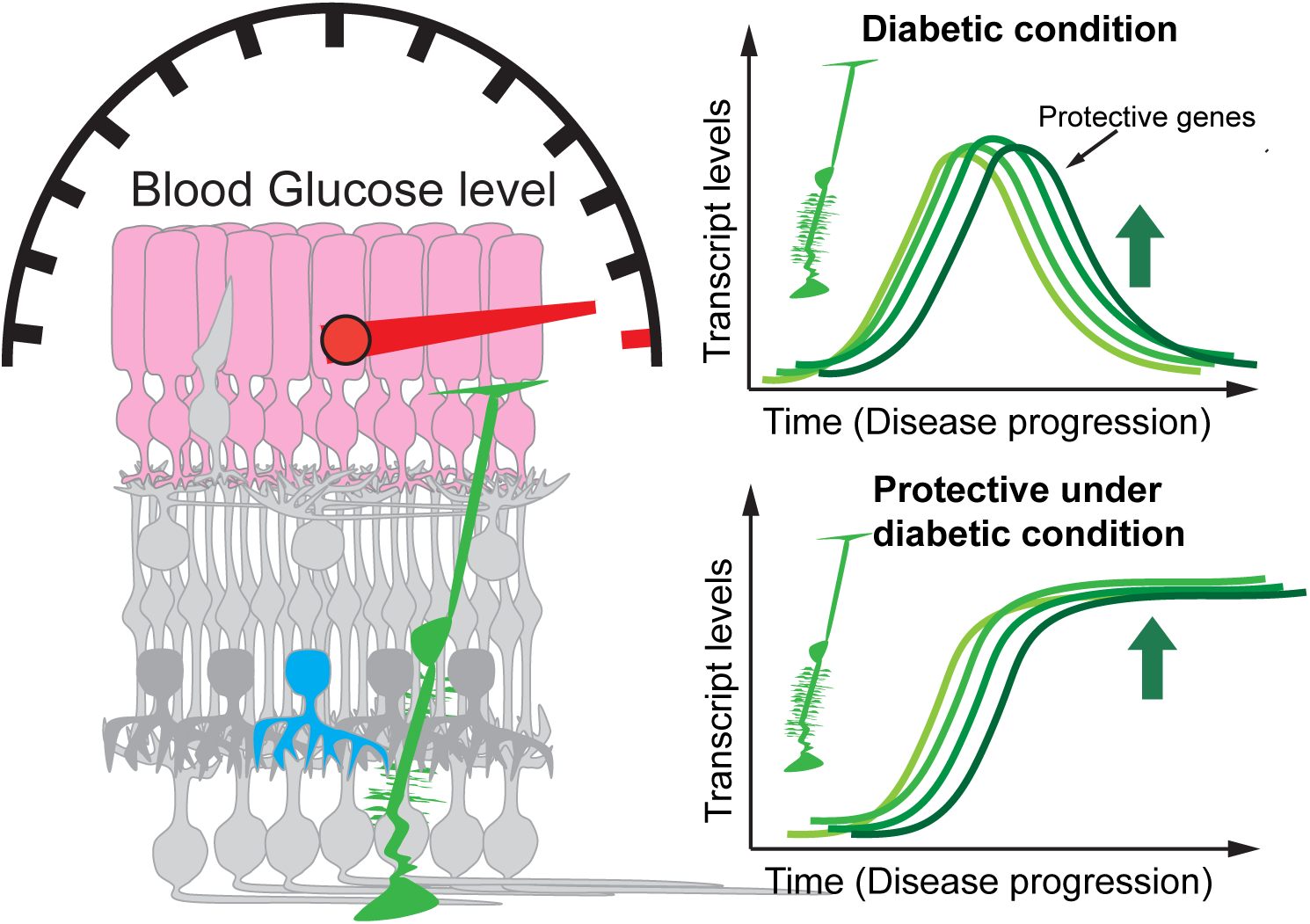
The working model of this study. Rod photoreceptors, A17 amacrine cells, and Müller glial cells exhibit transcriptional sensitivity to diabetes during the early stages of DR, as revealed by scRNA-seq. Notably, Müller glial cells initiate the early upregulation of protective genes, acting as a defense mechanism against diabetes-induced damage in the retina. However, this adaptive response wanes as the disease progresses. Elevating the expression of the genes transiently upregulated in Müller glial cells may effectively shield the retina from diabetes-induced damage.

Beyond the well-known impact of diabetes on retinal vasculature, various cell types within the neural retina exhibit susceptibility to diabetes, with some contributing to the early development of DR^6^. Our study underscores the potential roles of rod photoreceptor cells, A17 amacrine cells, and Müller glial cells in DR.

Although extensive photoreceptor degeneration is not commonly reported in DR patients and animal models, the functionality of photoreceptors is affected. Given their status as a major source of reactive oxygen species, photoreceptors can also contribute to DR development. The loss of photoreceptors can mitigate DR severity, possibly by reducing hypoxia^16^. In alignment with these findings, genes involved in oxidative phosphorylation and aerobic respiration were significantly upregulated in rods in our scRNA-seq dataset at both 1 month and 2 months post STZ injection. At 3 months post-STZ injection, genes linked to these pathways showed no significant alterations. Notably, there are fewer upregulated DEGs in the rod cluster at 3 months post-STZ compared to 1 or 2 months post-STZ (Figure 1D). Analysis of these upregulated DEGs did not reveal any enriched GO terms. These results imply a possible overall attenuation of rod responses at the transcript level as the disease progresses, warranting further investigation. Importantly, our study provides a candidate gene list for the observed rod changes in the literature, offering novel avenues for future testing and potential enhancement of therapeutic strategy design.

Neuronal dysfunction in the inner retina is also a recognized phenomenon in early DR models and human patients^36^. Oscillatory potentials (OPs), indicative of interactions between retinal bipolar and amacrine cells^37,38^, consistently undergo alterations during the early stages of DR in both animal models and humans^39^. Diabetic rodent models have reported reduced OP amplitude and delayed OP timing, indicating the induction of dysfunction in retinal amacrine cells in the early stages of diabetes^40,41^. Among various amacrine cell types, studies highlight the dysfunction of AII and A17 amacrine cells in response to diabetes, such as a diminished Ca^2+^ response to glutamate in A17 amacrine cells^42,43^. Our scRNA-seq data revealed a distinct cluster with gene expression profiles akin to A17 amacrine cells, exhibiting significant transcript-level changes at 1 month after STZ. The upregulated DEGs in this cluster are prominently associated with regulating cellular responses to metal ions and cytokine production. Further exploration is warranted to unveil the underlying molecular mechanisms, investigating whether A17 amacrine cells also actively contribute to DR development, as observed in photoreceptor cells.

Another cell type sensitive to diabetes in our scRNA-seq dataset is Müller glial cells, which can exert both beneficial and detrimental influences on DR development. On the positive side, the conditional ablation of Müller glial cells in mature retinas mirrors advanced-stage DR, manifesting as neurodegeneration, vascular leakage, and neovascularization ^20,44,45^. Notably, manipulating specific factors in Müller glial cells, such as αA-crystallin that regulates the ER stress pathway, has shown promise in alleviating DR development^46–50^. Conversely, studies indicate that Müller glial cells serve as critical sources of pro-inflammatory and neurotoxic factors, contributing to diabetes-induced inflammatory responses^32^. Nevertheless, the molecular mechanisms governing these responses remain incompletely elucidated, particularly the dynamic regulation of protective or detrimental genes in Müller glial cells as the disease progresses. Our study unveiled that Müller glial cells predominantly upregulate genes with protective roles at 1 month after STZ, yet this beneficial effect diminishes as the disease advances. Instead, genes related to oxidative phosphorylation or blood coagulation were upregulated at 2 or 3 months after STZ, respectively. Based on these findings, enhancing the activity of protective genes/pathways while inhibiting potential detrimental genes/pathways could represent an effective therapeutic strategy for future DR treatment, aiming to shield the retina from diabetes-induced damage.

Taken together, our scRNA-seq dataset highlights the potential susceptibility of rod, A17 amacrine cells, and Müller glial cells to diabetes. Given the depth of the method, our analysis focused on cell types or subtypes with a substantial number of captured cells. Notably, certain cell types known to contribute to DR, such as microglia, retinal ganglion cells, and retinal endothelial cells, were omitted due to low capture rates. Enhancing our understanding necessitates further investigation, involving an increase in cell numbers or targeted enrichment of specific cell types to provide a comprehensive view of diabetes-induced, cell-type-specific responses in the retina.

Furthermore, we identified differentially expressed gene sets across various retinal cell types, offering valuable molecular insights into the cell-type-specific responses triggered by diabetes. Despite the discussed limitations of the scRNA-seq technique, we successfully identified genes previously known to be dysregulated in the retina in response to diabetes. For example, we observed significant upregulation of *Fgf2* in rods and *Gfap* in Müller glial cells, aligning with published literature and attesting to the robustness of our assay. More importantly, our analyses revealed novel candidate genes with potential roles in protecting the retina from diabetes-induced damage. We specifically focused on dissecting the role of Zfp36 in Müller glial cells during DR, and revealed its ability to maintain Müller glial quiescence, inhibit gliosis, and alleviate DR-related phenotypes. Regrettably, determining whether Zfp36 can exert an interventive role is currently hindered by technical challenges. The intravitreal injection of AAVs proves intricate in adult rats post-STZ injection, with the added limitation of infecting only a small region of the retina. Future enhancements to the injection method in adult rats could overcome this limitation. Moreover, pinpointing the precise targets of Zfp36 proved challenging, given the difficulties encountered in obtaining enough Müller glial cells with either depleted or over-expressed *Zfp36*. Despite Zfp36’s recognized role as a potent inflammation inhibitor, we did not observe noteworthy upregulation of inflammation-related genes or pathways three months after STZ injection, a period characterized by reduced Zfp36 expression. It is conceivable that the impact of Zfp36 depletion requires a timeframe longer than three months to manifest, or alternatively, Zfp36 may exert its effects by regulating additional pathways, such as those associated with oxidative stress genes, which exhibit significant upregulation three months post STZ injection. Nevertheless, while the precise targets of Zfp36 in Müller glial cells await clarification, this study has established a framework for exploring other genes expressed in Müller glia that may harbor potential protective or detrimental effects in rat models in vivo. In fact, beyond Zfp36, we identified transient upregulation of additional genes in Müller glial cells during the early stages of DR, encompassing anti-inflammatory, anti-oxidative stress, anti-apoptotic, and anti-prefoliation pathways (Figure 2C and S3). Investigating whether the multiplex activation of these protective genes can more efficiently shield the retina from diabetes-induced damage can be an important avenue for future studies.

In summary, our scRNA-seq analysis delineated diabetes-induced transcriptomic changes in the rat retina at various stages following diabetes induction, pinpointing retinal cell types particularly responsive to diabetes. Of these diabetes-sensitive cell types, Müller glial cells exhibited rapid transcript-level responses to diabetes, transiently upregulating potential protective genes. Subsequent targeted investigations demonstrated that the overexpression of *Zfp36*, one of the identified protective gene candidates, could safeguard the retina from diabetes-induced damage in diabetic rat models. Enhancing intrinsic protective pathways emerges as a prospective therapeutic strategy for DR. Furthermore, the homogeneous nature of the rat models facilitated the discovery of gene expression changes at a high resolution, a challenging feat in human samples. Future validation of the dynamic regulation of protective genes in Müller glial cells in response to diabetes in human NPDR samples holds promise for a deeper comprehension of DR progression and the exploration of innovative therapeutic approaches for DR.

## Material and Methods

### Animals

Our study examined male rats because rats have gender sensitivity to streptozotocin (STZ). Female rats appear to be largely resistant to the effects of low-dose STZ. We obtained wild-type rats (SAS Sprague Dawley, stock #400) from Charles River Laboratories. Approval for all experiments involving laboratory animals was granted by the Administrative Panel on Laboratory Animal Care (APLAC) at Stanford University School of Medicine. Throughout the study, all rats were maintained under specific pathogen-free conditions at the animal facility of Stanford University School of Medicine.

### Induction of Diabetes

STZ injection was carried out in accordance with the established protocol^51^. On the injection day, 8-week-old male rats underwent a 6-8 hour fasting period. Citrate buffer (0.1M Sodium Citrate, pH 4.5) and a 10% sucrose (w/v) solution were freshly prepared. STZ (Sigma #0130) was dissolved in citrate buffer, reaching a final concentration of 10mg/ml immediately prior to injection. The intraperitoneal route was employed for injection, with a dosage of 65mg/kg rat. Fasting glucose levels were monitored at 2, 7 days, 2 weeks, 1-, 2-, or 3-months post-injection. Male rats exhibiting consistently elevated fasting glucose levels (measured by the One Touch Ultramini Glucose monitoring system) exceeding 250mg/dl at all time points were included in the study. Insulin was intentionally omitted to mitigate confounding factors.

### Adeno-Associated Virus (AAV) Production and Delivery

The AAV-pGFAP-EGFP backbone plasmid, generously provided by Dr. Wenjun Xiong (City College of Hong Kong), served as the foundation for constructing all AAV plasmids using restrictive enzyme-based cloning. The shRNA cassettes, designed to target rat *Zfp36*, were crafted in accordance with established methodologies^29^. The AAV-pGFAP-EGFP-LacZ-shRNA (AAV-Ctrl) and AAV-pGFAP-EGFP-Zfp36-shRNA (AAV-Zfp36-KD) plasmids were generated by integrating the LacZ-shRNA or Zfp36-shRNA cassette (shRNA-76, Figure S8) into the 3’UTR of the *Egfp* gene, following a published study^29^. The AAV-pGFAP-ZR-p2A-EGFP plasmid was constructed by inserting a codon-optimized rat Zfp36 ORF, designed to be impervious to Zfp36-shRNA recognition, into the AAV-pGFAP-EGFP plasmid. The AAV-pGFAP-Zfp36-p2A-EGFP plasmid was created by introducing the wildtype *Zfp36* ORF into the AAV-pGFAP-EGFP plasmid backbone. Details of primers and sequences are outlined in Table S1.

Recombinant AAV7m8 viruses, produced as detailed in a prior publication^33^, underwent titer determination through SDS-PAGE gels. AAV intravitreal injection was administered to P4 neonatal rats using a pulled glass needle controlled by the Femtojet microinjection system (Eppendorf), as previously described^33^. Approximately 1.5 µl of AAV7m8 (∼10^12 gc/ml) was delivered into the intravitreal space.

### Histology and Immunohistochemistry

Rat eyeballs were fixed in 4% paraformaldehyde (PFA) in 1×PBS (pH 7.4) for 2 hours at room temperature. Subsequently, retinas were dissected in 1xPBS and equilibrated at room temperature in a sequential sucrose gradient (5% sucrose in 1× PBS, 5 min; 15% sucrose in 1× PBS, 15 min; 30% sucrose in 1× PBS, 1 h; 1:1 mixed solution of OCT and 30% sucrose in PBS, 4°C, overnight), followed by embedding in OCT and storage at −80°C. Cryosections of 20 μm thickness were prepared using a Leica CM3050S cryostat (Leica Microsystems). These retinal cryosections were briefly washed in 1× PBS, incubated in 1× PBS with 0.2% Triton for 20 min, and subsequently blocked for 30 min in a blocking solution (0.1% Triton, 1% BSA, and 10% donkey serum in 1x PBS). Slides were incubated with primary antibodies diluted in blocking solution in a humidified chamber at 4°C overnight. Following three washes in 1× PBS with 0.1% Triton, slides underwent a 1-hour incubation with secondary antibodies and DAPI (Sigma-Aldrich; D9542). After three washes with 1× PBS with 0.1% Triton, the slides were mounted in Fluoromount-G (Southern Biotechnology Associates).

Primary antibodies utilized include chicken anti-GFP (Abcam, ab13970, 1:1000), rabbit anti-Sox9 (Abcam, ab185966, 1:500), chicken anti-GFAP (Fisher scientific, NBP105198, 1:500), rabbit anti-Pax6 (Thermo Fisher Scientific, 42-6600, 1:500), sheep anti Chx10(Vsx2) (Exalpha Biologicals, X1179P, 1:100), Guinea pig anti RBPMS (PhosphoSolutions, 1832-RBPMS, 1:500), and goat anti Iba1 (Abcam, ab5076, 1:500).

### mRNA fluorescence in situ Hybridization

Rat retinal tissues were promptly dissociated following established procedures^52^. Post-dissociation, retinal cells were incubated on Poly-D-Lysine (PDL, 0.1mg/ml, Millipore) treated round coverslips (Fisher Scientific, 50-121-5159) for 45 minutes at 37°C. Subsequently, the cells were fixed on the coverslips with 4% paraformaldehyde (PFA) in 1×PBS (pH 7.4) for 30 minutes at room temperature. Following fixation, the cells underwent dehydration through a gradient of 50%, 70%, and 100% clean ethanol and were stored in 100% ethanol at -20°C. Probes for mRNA in situ hybridization were purchased from ACDBio, and mRNA FISH was conducted according to commercial protocols. FISH signals were captured using Zeiss confocal LSM 880 and Olympus FV3000 confocal microscopies. Image analysis was carried out using Zen software (Zeiss), and quantification was performed using Fiji (NIH).

### Western blot

Freshly dissected rat retinas were placed in RIPA buffer (Radioimmunoprecipitation assay buffer, Abcam #ab156034) and heated at 95°C for 10 minutes. Protein concentrations were determined using the BCA protein kit (Pierce™ BCA Protein Assay Kit #23228). Equal protein amounts were loaded onto a 4%-20% SDS-PAGE gel (BIO-RAD, #4561094) and electrophoresed at 130V and 30A for 1.5 hours. The gel was then transferred to a 0.2mm PVDF membrane (BIO-RAD, #1704156) using a semi-dry transfer system (BIO-RAD, 1704150EDU). Following blocking with a 5% non-fat milk solution, the membrane was incubated with primary antibodies at 4°C overnight, starting with rabbit anti-Zfp36 (Fisher Scientific, PIPA540876, 1:500). After a 30-minute wash with 1X TBST (20mM Tris, 150mM NaCl, 0.1% Tween20), the membrane was incubated with a horseradish peroxidase-conjugated secondary antibody (Amersham, NA934-100U, 1:5000) at room temperature for 2 hours. Subsequent washes were followed by incubation with the SuperSignal West Pico PLUS chemiluminescent substrate mix (Thermo Scientific, #34580). Chemiluminescent signals were captured using the Amersham ImageQuant 600 (Cytiva) imaging system. After imaging, Zfp36 signals were stripped from the membrane using western blot stripping buffer (ThermoFisher, 21059), and rabbit anti-GAPDH (Abcam, ab9485, 1:1000) was applied. GAPDH signals were obtained using the same procedure as described for Zfp36.

### Retina Vessel Permeability Assay and whole mount antibody staining

Rats were deeply anesthetized through ketamine injection and inhalation of isoflurane. EZ-Link™ Sulfo-NHS-LC-Biotin (Thermo Fisher Scientific; 21335) at 0.4 mg/g body weight was injected into the left ventricle using a 26-gauge needle and a 1 ml syringe. Rats exhibiting a continuous, steady heartbeat for 7 minutes were included in the study, while those with a cessation of heartbeat within this timeframe were excluded from subsequent analyses. After 15 minutes of tracer circulation, eyeballs were enucleated and fixed in 4% paraformaldehyde (PFA) in 1X PBS (pH 7.4) for 1 hour at room temperature, followed by overnight fixation at 4 °C. Retinas were then dissected and immersed in a wholemount blocking solution (3% Triton X-100, 0.5% Tween 20, 1% bovine serum albumin, 0.1% sodium azide, and 10% donkey serum in 1X PBS) for 24 hours at 4°C before signal detection.

For NHS-Biotin detection, retinas were incubated with Alexa Fluor 555-conjugated Streptavidin (ThermoFisher, S21381, 1:1000) for 2 hours at room temperature and 24 hours at 4°C. For IB4 staining, retinas were incubated with Alexa Fluor 647-conjugated Isolectin GS-IB4 (ThermoFisher, I32450, 1:1000). For desmin staining, retinas were incubated with desmin antibody (Rabbit anti-Desmin, Abcam, ab15200, 1:500) in wholemount blocking solution at room temperature for 4 hours and at 4°C for 48 hours. Subsequent steps included incubation with fluorochrome-conjugated secondary antibodies at room temperature for 2 hours and 4°C for 24 hours, followed by mounting using Fluoromount-G.

### Single Cell RNA sequencing (scRNA-seq) and data analysis

Rat retinal tissues were dissociated following a previously established protocol^52^. To selectively deplete rod photoreceptors, mouse anti-rat CD73 (Fisher Scientific, BDB551123, 5 μl per 10^7 cells) and mouse IgG microbeads (Miltenyi Biotech, 130-047-102, 10 μl per 10^7 cells) were sequentially introduced into the dissociated cell suspension and incubated at room temperature for 10 minutes with each reagent. The cell suspension was then processed through an LD column (Miltenyi Biotec, 130-042-901), and the follow-through was collected, followed by centrifugation at 300 rcf for 5 minutes. To eliminate dead cells, the resulting pellet was resuspended in 100 μl of microbeads from the dead cell removal kit (Miltenyi Biotec, 130-090-101) and processed according to the commercial protocol. The cells were ultimately resuspended in 100 μl of DMEM + 0.4% BSA (Sigma-Aldrich, A1595-50ml) and loaded onto a 10x Genomics Chromium controller. Single-cell RNA-Seq libraries were prepared using the Chromium Gene Expression 3’ kit v3.1 (PN-1000269). Library quality was assessed using a 2100 Bioanalyzer (Agilent) with a high-sensitivity DNA kit (Agilent, #5067-4626). Illumina HiSeq4000 platform (2 x 150 paired end) was used for sequencing.

Single-cell RNA expression matrices were generated by aligning sequencing reads to a pre-built rat reference (mRatBN7.2) using the Cell Ranger (v.6.1.1) pipeline and subsequently loaded into R (v.4.1) for downstream analyses. DoubletFinder (McGinnis et al., 2019) was employed to remove doublets. Filtered matrices underwent Seurat (v.4.1) processing for quality control, clustering, and visualization using standard workflows. In brief, genes detected in fewer than 3 cells and cells with fewer than 600 genes or more than 25% reads as mitochondrial were excluded from analyses. Samples were normalized and integrated using 2000 variable genes (FindIntegrationAnchors and IntegrateData). Clusters were generated based on the top 30 principal components (PCs) (FindNeighbors function and FindClusters function, resolution = 0.5) and visualized by Uniform Manifold Approximation and Projection (UMAP). For identification of conserved markers in each cluster, differential gene expression tests were performed on each cluster versus all other clusters (FindConservedMarkers). Clusters were categorized into 10 major cell types based on their conserved markers. Differentially expressed genes (DEGs) were identified using a Wilcoxon rank-sum test (FindAllMarkers) between Ctrl and STZ in each cell cluster. DEGs with |log2 (fold change)| > 1 and p-value < 0.05 were selected for input into GO enrichment analyses. To query MG or cluster 16 on published reference datasets^23,24^, the Unimodal UMAP projection function (Mapquery, Seurat 4.0.6) was employed.

### Imaging and analysis

Retinal images were captured using a Zeiss LSM880 inverted confocal microscope. The images presented in Figures 3, 4, and 5 represent maximum projections of 5-10μm tissue sections, and their quantification was performed using Fiji software.

## QUANTIFICATION AND STATISTICAL ANALYSIS

Statistical analyses were conducted according to the specifications outlined in the figure legends. GraphPad Prism v.9 (GraphPad) facilitated the execution of statistical procedures, with graphs and error bars reflecting means ± SD. The study utilized an unpaired t-test with Welch’s correction (two-tailed). Statistical significance was defined as a P value <0.05, denoted by asterisks: * for P < 0.05, ** for P < 0.01, *** for P < 0.001, and **** for P < 0.0001, while "ns" indicated P ≥ 0.05. N number is indicated in figures.

### Data availability

Data has been uploaded to the GEO database awaiting approval.

## Acknowledgements

The members of the Wang laboratory provided valuable discussion and support for this project.

## Author Contribution

S.W. conceived and supervised the study. S.W. and C.H.L. designed the experiments. Bogdan and S.W. performed the bioinformatic analyses. C.H.L., M.R.W., A.E.D., L.L., Y.S. and A.X. performed the experiments. S.W. wrote the manuscript. F.P. edited the manuscript. All authors provided critical feedback.

## Funding and additional information

Support was provided by American Diabetes Association (1-16-INI-16 to S.W.), NEI 1R01EY03258501 (to S.W.), 1R01EY03379201 (to S.W.) and NIH P30EY026877 to Stanford Ophthalmology, and by NEI 1R01EY033527 (to P.E.F.) and NIH P30EY007003 to the University of Michigan, as well as Eversight and Research to Prevent Blindness (to P.E.F.). The content is solely the responsibility of the authors and does not necessarily represent the official views of the National Institutes of Health.

