## Supplementary Figures and Table for "Induction of a Müller glial-specific protective pathway safeguards the retina from diabetes induced damage"

**Supplementary information**

1. **Figure S1 to S8 with figure legends**
2. **Supplementary table S1**

Figure S1

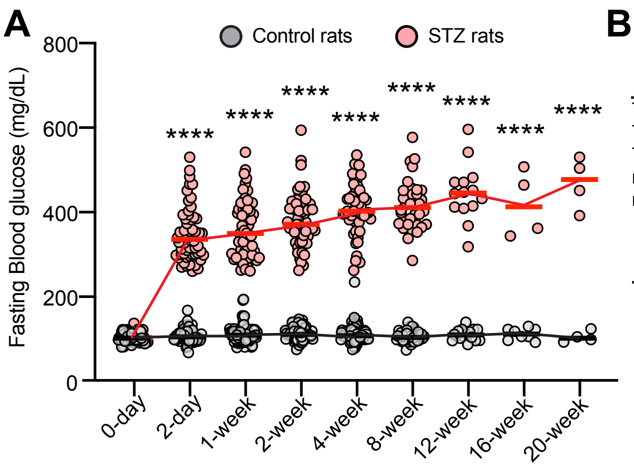


B

**Before QC**

| | 1mWT | 1mSTZ | 2mWT | 2mSTZ | 3mWT | 3mSTZ |
|---|---|---|---|---|---|---|
| Total Cells | 9206 | 9500 | 8946 | 13749 | 13996 | 15088 |
| Total reads | 160,710,986 | 154,626,701 | 211,362,951 | 240,559,450 | 162,235,045 | 156,257,836 |
| Total genes | 25399 | 25399 | 25399 | 25399 | 25399 | 25399 |
| median genes/cell | 1786 | 1617 | 1897 | 1479 | 1517 | 1409 |
| median UMI counts/cells | 3253 | 2843 | 3816 | 2575 | 2544 | 2376 |

**After QC**

| | 1mWT | 1mSTZ | 2mWT | 2mSTZ | 3mWT | 3mSTZ |
|---|---|---|---|---|---|---|
| Total cells | 7335 | 8385 | 7868 | 11725 | 10998 | 13759 |
| Total genes | 17419 | 17574 | 17552 | 17840 | 17614 | 17864 |
| Median genes/cell | 1956 | 1736 | 1969 | 1579 | 1686 | 1484 |
| Median counts/cells | 3680 | 3148 | 4037 | 2781 | 2972 | 2552 |

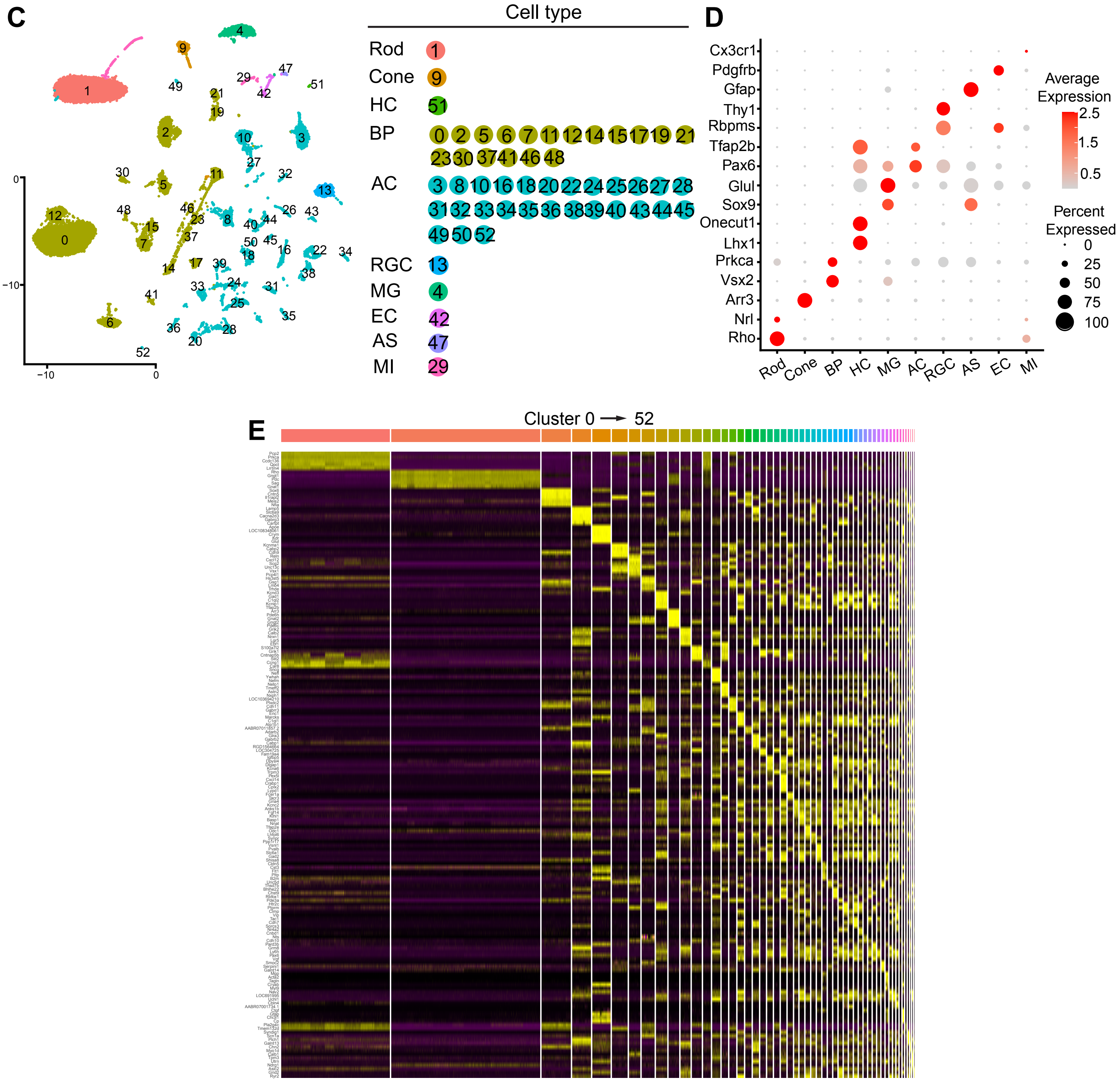

**Figure S1**. **The quality control analyses of scRNA-seq data (related to Figure 1)**
**(A)** Fasting blood glucose levels in male rats were monitored at 2 days, 1, 2, 4, 8, 12, 16, and 20 weeks post citrate buffer (control rats) or STZ (STZ rats) injection. Mean ± SD. Unpaired t-test with Welch's correction (two-tailed). **** $P < 0.0001$. N numbers are indicated in the charts. **(B)** Summary of quality control metrics for each sample. **(C)** The relationship between cluster number and cell type. **(D)** Markers employed for the identification of major retinal cell types. **(E)** The top 5 differentially expressed genes for each cluster.

Figure S2

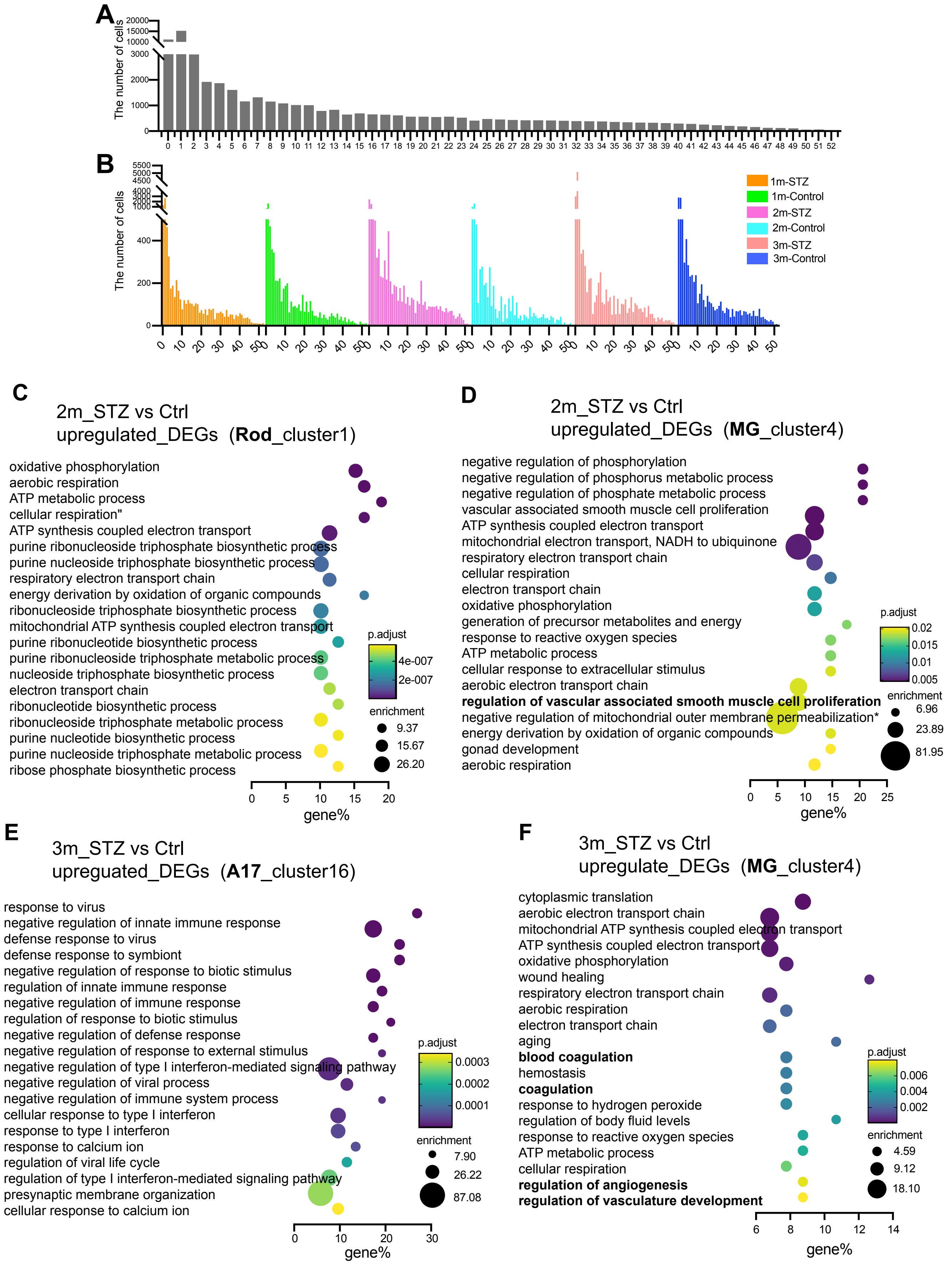

A
The number of cells
20000
15000
10000
3000
2000
1000
0
0 1 2 3 4 5 6 7 8 9 10 11 12 13 14 15 16 17 18 19 20 21 22 23 24 25 26 27 28 29 30 31 32 33 34 35 36 37 38 39 40 41 42 43 44 45 46 47 48 49 50 51 52
B
The number of cells
5500
5000
4500
4000
400
200
0
1m-STZ
1m-Control
2m-STZ
2m-Control
3m-STZ
3m-Control
0 10 20 30 40 50
C
2m_STZ vs Ctrl
upregulated_DEGs (Rod_cluster1)
oxidative phosphorylation
aerobic respiration
ATP metabolic process
cellular respiration"
ATP synthesis coupled electron transport
purine ribonucleoside triphosphate biosynthetic process
purine nucleoside triphosphate biosynthetic process
respiratory electron transport chain
energy derivation by oxidation of organic compounds
ribonucleoside triphosphate biosynthetic process
mitochondrial ATP synthesis coupled electron transport
purine ribonucleotide biosynthetic process
purine ribonucleoside triphosphate metabolic process
nucleoside triphosphate biosynthetic process
electron transport chain
ribonucleotide biosynthetic process
ribonucleoside triphosphate metabolic process
purine nucleotide biosynthetic process
purine nucleoside triphosphate metabolic process
ribose phosphate biosynthetic process
p.adjust
4e-007
2e-007
enrichment
9.37
15.67
26.20
0 5 10 15 20
gene%
D
2m_STZ vs Ctrl
upregulated_DEGs (MG_cluster4)
negative regulation of phosphorylation
negative regulation of phosphorus metabolic process
negative regulation of phosphate metabolic process
vascular associated smooth muscle cell proliferation
ATP synthesis coupled electron transport
mitochondrial electron transport, NADH to ubiquinone
respiratory electron transport chain
cellular respiration
electron transport chain
oxidative phosphorylation
generation of precursor metabolites and energy
response to reactive oxygen species
ATP metabolic process
cellular response to extracellular stimulus
aerobic electron transport chain
regulation of vascular associated smooth muscle cell proliferation
negative regulation of mitochondrial outer membrane permeabilization*
energy derivation by oxidation of organic compounds
gonad development
aerobic respiration
p.adjust
0.02
0.015
0.01
0.005
enrichment
6.96
23.89
81.95
0 5 10 15 20 25
gene%
E
3m_STZ vs Ctrl
upreguated_DEGs (A17_cluster16)
response to virus
negative regulation of innate immune response
defense response to virus
defense response to symbiont
negative regulation of response to biotic stimulus
regulation of innate immune response
negative regulation of immune response
regulation of response to biotic stimulus
negative regulation of defense response
negative regulation of response to external stimulus
negative regulation of type I interferon-mediated signaling pathway
negative regulation of viral process
negative regulation of immune system process
cellular response to type I interferon
response to type I interferon
response to calcium ion
regulation of viral life cycle
regulation of type I interferon-mediated signaling pathway
presynaptic membrane organization
cellular response to calcium ion
p.adjust
0.0003
0.0002
0.0001
enrichment
7.90
26.22
87.08
0 10 20 30
gene%
F
3m_STZ vs Ctrl
upregulate_DEGs (MG_cluster4)
cytoplasmic translation
aerobic electron transport chain
mitochondrial ATP synthesis coupled electron transport
ATP synthesis coupled electron transport
oxidative phosphorylation
wound healing
respiratory electron transport chain
aerobic respiration
electron transport chain
aging
blood coagulation
hemostasis
coagulation
response to hydrogen peroxide
regulation of body fluid levels
response to reactive oxygen species
ATP metabolic process
cellular respiration
regulation of angiogenesis
regulation of vasculature development
p.adjust
0.006
0.004
0.002
enrichment
4.59
9.12
18.10
6 8 10 12 14
gene%

**Figure S2. The number of cells captured in each cluster, and the enriched Gene Ontology (GO) terms for rod, Müller glial cells, and A17 amacrine cells at 2- or 3-month post STZ injection (related Figure 1 and 2).**

**(A-B)** The quantity of retinal cells in each cluster post quality control is depicted, illustrating the total number of cells per cluster (A) and per cluster per condition (B). **(C-D)** The top 20 enriched GO terms for upregulated DEGs in rod (cluster 1, C) or Müller glial cells (cluster 4, D) at 2 months after STZ injection. **(E-F)** The top 20 enriched GO terms for upregulated DEGs in A17 amacrine cells (cluster 16, E) or Müller glial cells (cluster 4, F) at 3 months after STZ injection.

Figure S3

| Gene | Function | Role | Supplementary References |
|---|---|---|---|
| Stat3 | anti- or proinflammatory | Context-dependent | 53-55 |
| Cebpd | anti-apoptosis or pro-migration | Context-dependent | 56, 57 |
| Chka | oncogene | Negligent | 58 |
| Plat | activate microglia (act as a cytokine) | Negligent | 59 |
| Ctsc | pro-inflammation | Negligent | 60 |
| Slc3a2 | oncogene | Negligent | 61, 62 |
| Ano6 | oncogene | Negligent | 63 |
| Vav3 | anti-survival | Negligent | 64 |
| Ptprn | oncogene | Negligent | 65 |
| Plin2 | pro-inflammation | Negligent | 66 |
| Rnf19b | anti-inflammation | Protective | 67 |
| Lcn2 | Neuroprotective | Protective | 68 |
| Dclk1 | pro-survival | Protective | 69, 70 |
| Scg2 | anti-angiogenesis | Protective | 71, 72 |
| Enox1 | anti-apoptosis, anti-angiogenesis | Protective | 73, 74 |
| Slc7a11 | neuroprotective, anti oxidative stress | Protective | 75 |
| Ifrd1 | anti-inflammation, anti-apoptosis | Protective | 76 |
| Qsox1 | anti-inflammation | Protective | 77 |
| Zfp36 | anti-inflammation | Protective | 78 |
| Cyp26a1 | anti-apoptosis | Protective | 79-81 |
| Mt3 | anti-oxidative stress | Protective | 82, 83 |
| Mt1 | anti-oxidative stress | Protective | 84 |
| Akap12 | anti-oxidative stress, anti-inflammation, anti-angiogenesis | Protective | 85, 86 |
| AABR07040864.1 | anti-apoptosis | Protective | 87 |
| Cited2 | anti-inflammation | Protective | 88 |
| Mt2A | anti-oxidant, anti-apoptosis,detoxification, anti-proliferationanti-inflammation | Protective | 89 |
| Mt1m | anti-proliferation | Protective | 90 |
| Slc14a1 | neuroprotective | Protective | 91 |
| Cp | neuroprotective, anti-oxidative stress | Protective | 92 |
| Rnd3 | anti-proliferation | Protective | 93 |
| Dusp6 | neuroprotective | Protective | 94 |
| Timp1 | anti-inflammation | Protective | 95, 96 |
| Gdpd2 | Unknown | Unknown | |
| Ac128848.1 | Unknown | Unknown | |
| Sorcs1 | Unknown | Unknown | |
| Col5a3 | Unknown | Unknown | |

**Figure S3. The predicted roles of the DEGs that are temporarily upregulated at 1 month (up-down DEGs, 1m_STZ vs 1m_Ctrl) in Müller glial cells based on the literature (related to Figure 2)**

The gene name, function, predicted roles and references are listed.

### Figure S4

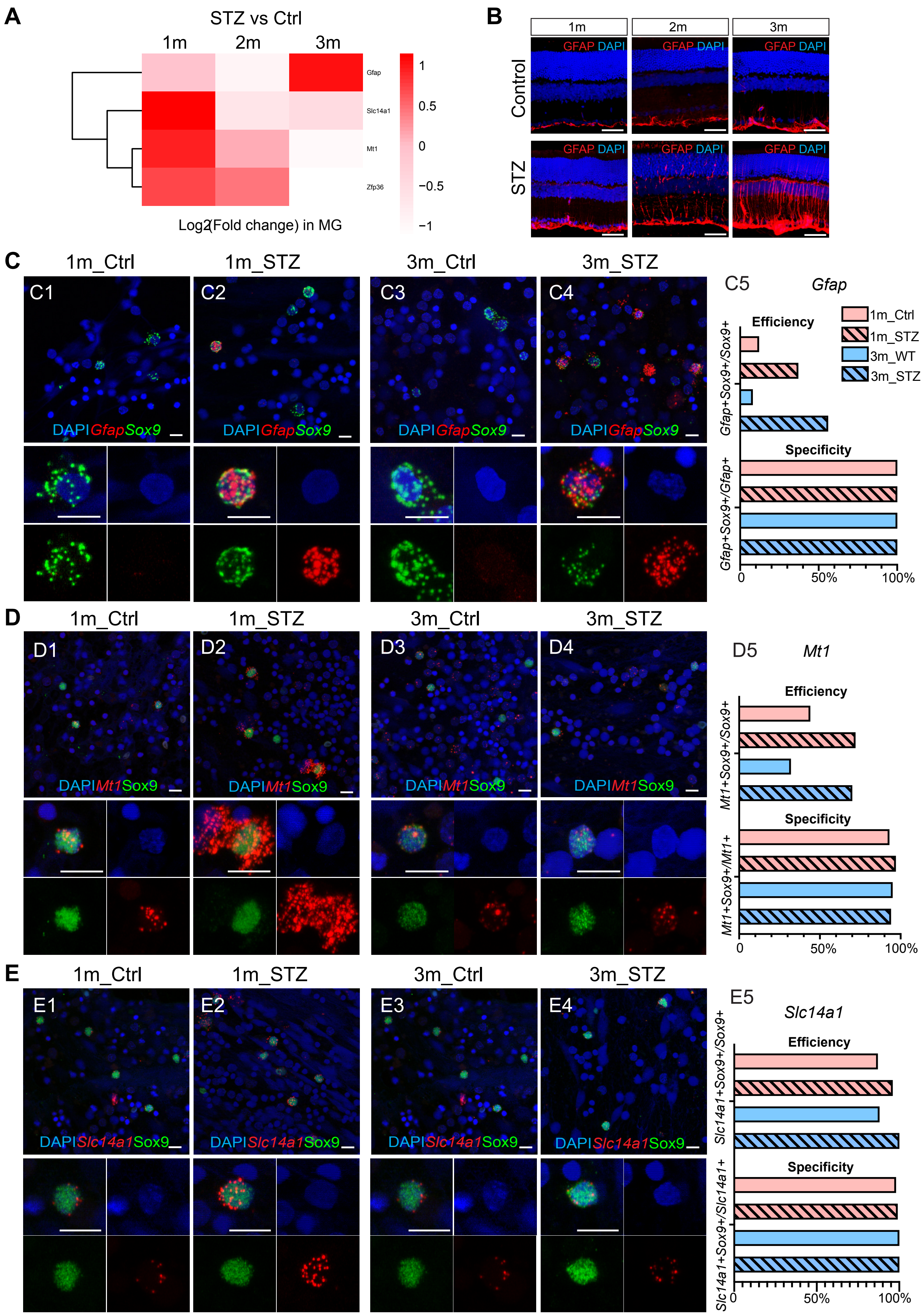
A
STZ vs Ctrl
1m
2m
3m
Gfap
Slc14a1
Mt1
Zfp36
1
0.5
0
−0.5
−1
Log2(Fold change) in MG
B
1m
2m
3m
Control
STZ
GFAP DAPI
C
1m_Ctrl
1m_STZ
3m_Ctrl
3m_STZ
C1
C2
C3
C4
DAPI Gfap Sox9
C5
Gfap
Efficiency
Specificity
1m_Ctrl
1m_STZ
3m_WT
3m_STZ
Gfap+Sox9+/Sox9+
Gfap+Sox9+/Gfap+
0
50%
100%
D
1m_Ctrl
1m_STZ
3m_Ctrl
3m_STZ
D1
D2
D3
D4
DAPI Mt1 Sox9
D5
Mt1
Efficiency
Specificity
Mt1+Sox9+/Sox9+
Mt1+Sox9+/Mt1+
0
50%
100%
E
1m_Ctrl
1m_STZ
3m_Ctrl
3m_STZ
E1
E2
E3
E4
DAPI Slc14a1 Sox9
E5
Slc14a1
Efficiency
Specificity
Slc14a1+Sox9+/Sox9+
Slc14a1+Sox9+/Slc14a1+
0
50%
100%

**Figure S4. The expression of *Gfap* and candidate protective genes in Müller glial cells at 1 to 3 months post citrate buffer or STZ injection (related to Figure 2)**

**(A)** Heatmap illustrating the $log_2$(fold changes) of *Gfap*, *Mt1*, *Slc14a1* and *Zfp36* in Müller glial cells (MG) at 1, 2, or 3 months after citrate buffer or STZ injection. **(B)** The expression of GFAP protein in rat retinas. 1m, 2m or 3m: 1, 2 or 3 months after injection. Control: with citrate buffer injection. STZ: with STZ injection. **(C)** *Gfap* expression (red signal). Sox9 smFISH signals (green) indicate Müller glial cells (MG). The percentage of MG expressing *Gfap* (efficiency) and the percentage of *Gfap*+ cells that are MG (specificity) are shown in A5. **(D)** *Mt1* expression (red signal). Sox9 antibody staining signals (green) indicate Müller glial cells (MG). The percentage of MG expressing *Mt1* (efficiency) and the percentage of *Mt1*+ cells that are MG (specificity) are shown in A5. **(E)** *Slc14a1* expression (red signal). Sox9 antibody staining signals (green) indicate Müller glial cells (MG). The percentage of MG expressing *Slc14a1* (efficiency) and the percentage of *Slc14a1*+ cells that are MG (specificity) are shown in A5.

### Figure S5

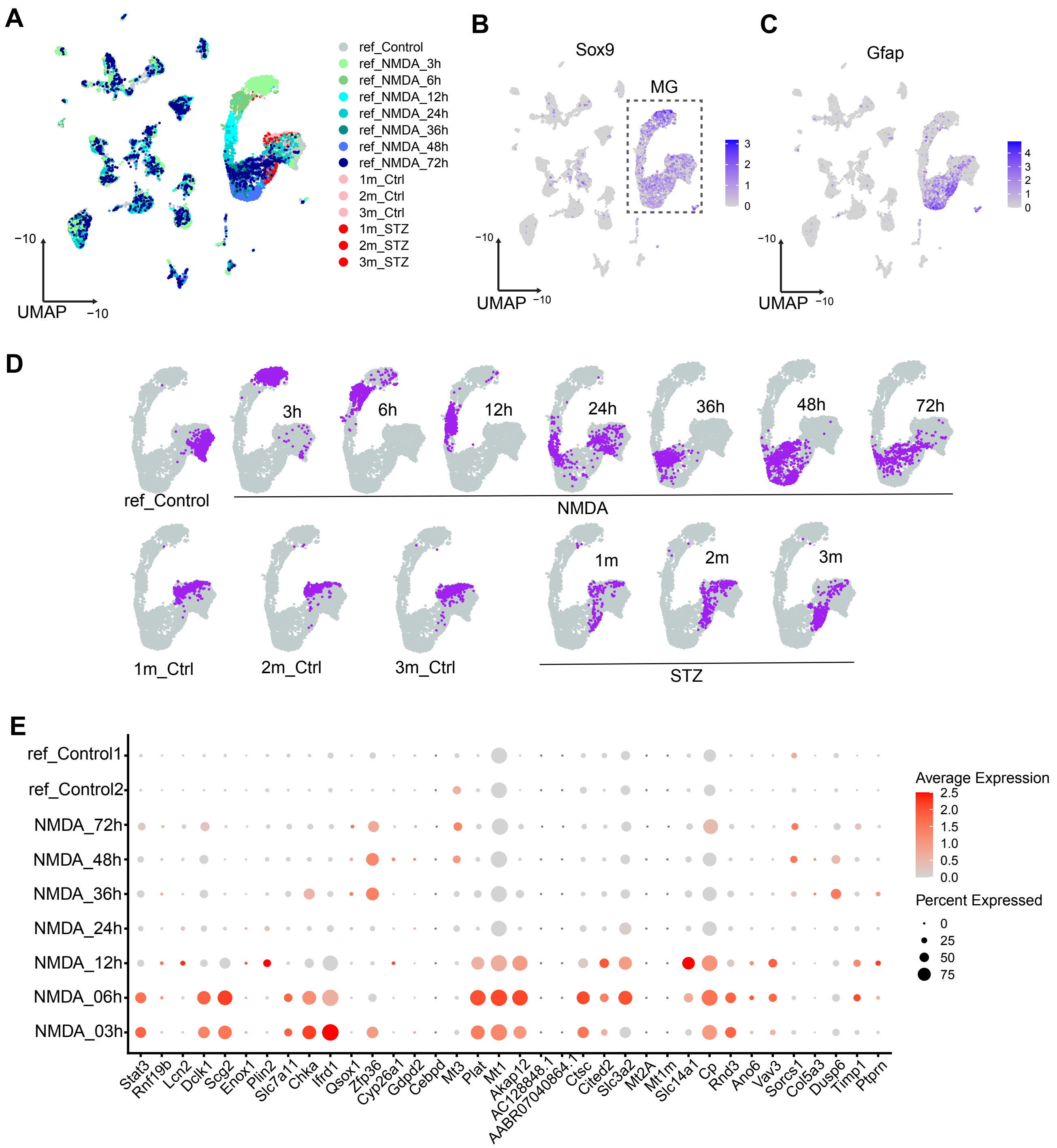

A
ref_Control
ref_NMDA_3h
ref_NMDA_6h
ref_NMDA_12h
ref_NMDA_24h
ref_NMDA_36h
ref_NMDA_48h
ref_NMDA_72h
1m_Ctrl
2m_Ctrl
3m_Ctrl
1m_STZ
2m_STZ
3m_STZ
UMAP
−10
−10
B
Sox9
MG
C
Gfap
D
ref_Control
3h
6h
12h
24h
36h
48h
72h
NMDA
1m_Ctrl
2m_Ctrl
3m_Ctrl
1m
2m
3m
STZ
E
ref_Control1
ref_Control2
NMDA_72h
NMDA_48h
NMDA_36h
NMDA_24h
NMDA_12h
NMDA_06h
NMDA_03h
Stat3
Rnf19b
Lcn2
Dclk1
Scg2
Enox1
Plin2
Slc7a11
Chka
Ifrd1
Qsox1
Zfp36
Cyp26a1
Gdpd2
Cebpd
Mt3
Plat
Mt1
Akap12
AC128848.1
AABR07040864.1
Ctsc
Cited2
Slc3a2
Mt2A
Mt1m
Slc14a1
Cp
Rnd3
Ano6
Vav3
Sorcs1
Col5a3
Dusp6
Timp1
Ptprn
Average Expression
Percent Expressed

**Figure S5.** A comparison between Müller glial cells (MG) from rat diabetic retinas and those from an acute injury mouse model, revealing partially overlapping transcriptional profiles.

**(A)** UMAP plot illustrating diabetic rat MG (query) and mouse MG subjected to NMDA treatment (reference). **(B)** UMAP plot illustrating the expression of *Sox9*, with the MG cluster outlined by a dotted line box. **(C)** UMAP plot illustrating the expression of *Gfap*, indicating gliosis in MG. **(D)** UMAP plots illustrating MG profiles under different conditions. Wild type control mouse MG are depicted on the top, while control or STZ rat MG are illustrated at the bottom. **(E)** Dotplot showing the expression of DEGs that are temporarily upregulated at 1 month (1m up-down DEGs, 1m_STZ vs 1m_Ctrl) in NMDA treated mouse MG at various time points post NMDA injection.

Figure S6

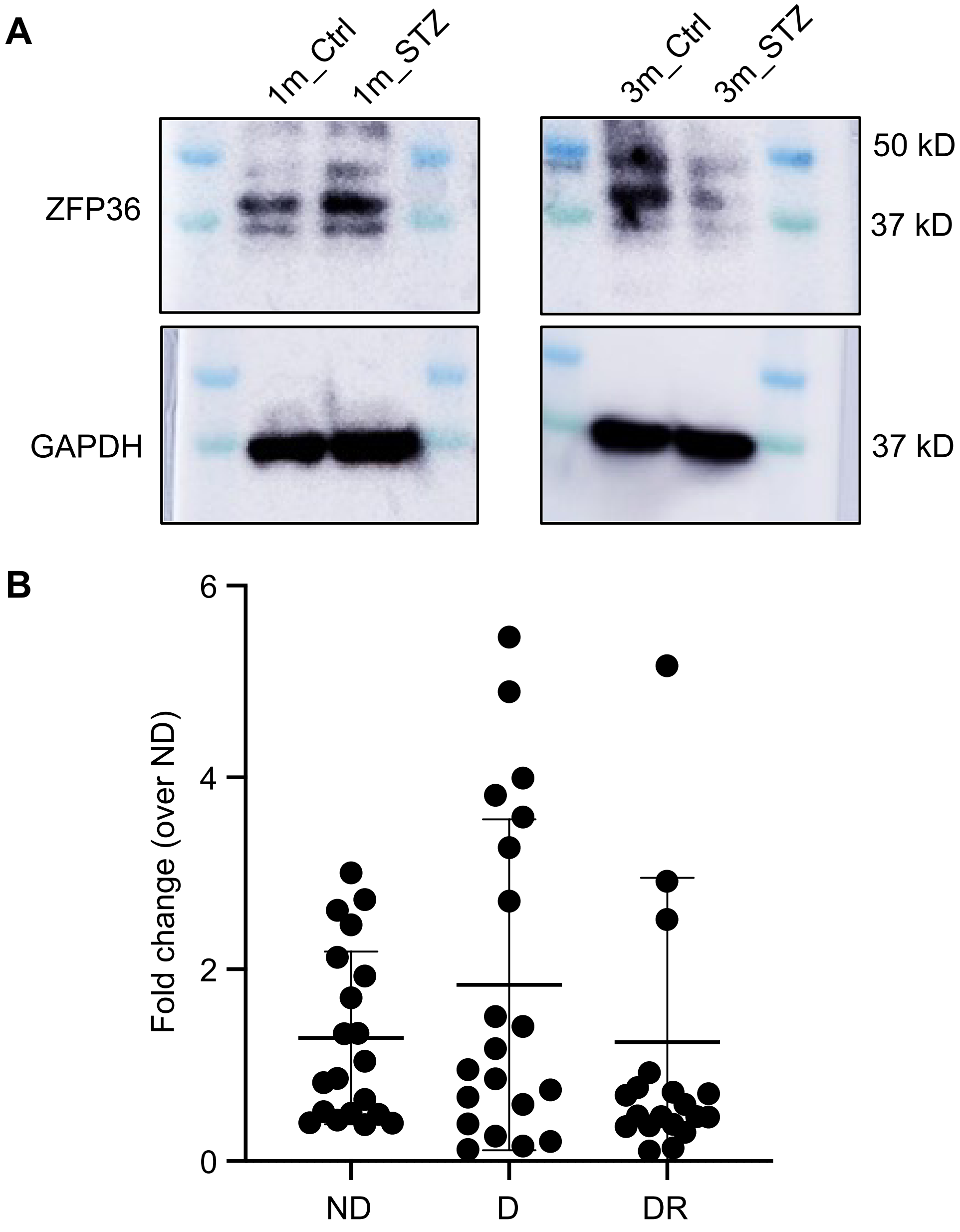


**Figure S6. Zfp36 is transiently upregulated in diabetic retinas (related to Figure 3). (A)** ZFP36 and GAPDH were detected by western blotting in rat retinas at 1 month or 3 months post STZ injection. **(B)** The expression of *Zfp36* in human retinas was detected by qPCR. ND: healthy control; D: diabetic control without DR; DR: human retina samples with DR.

Figure S7

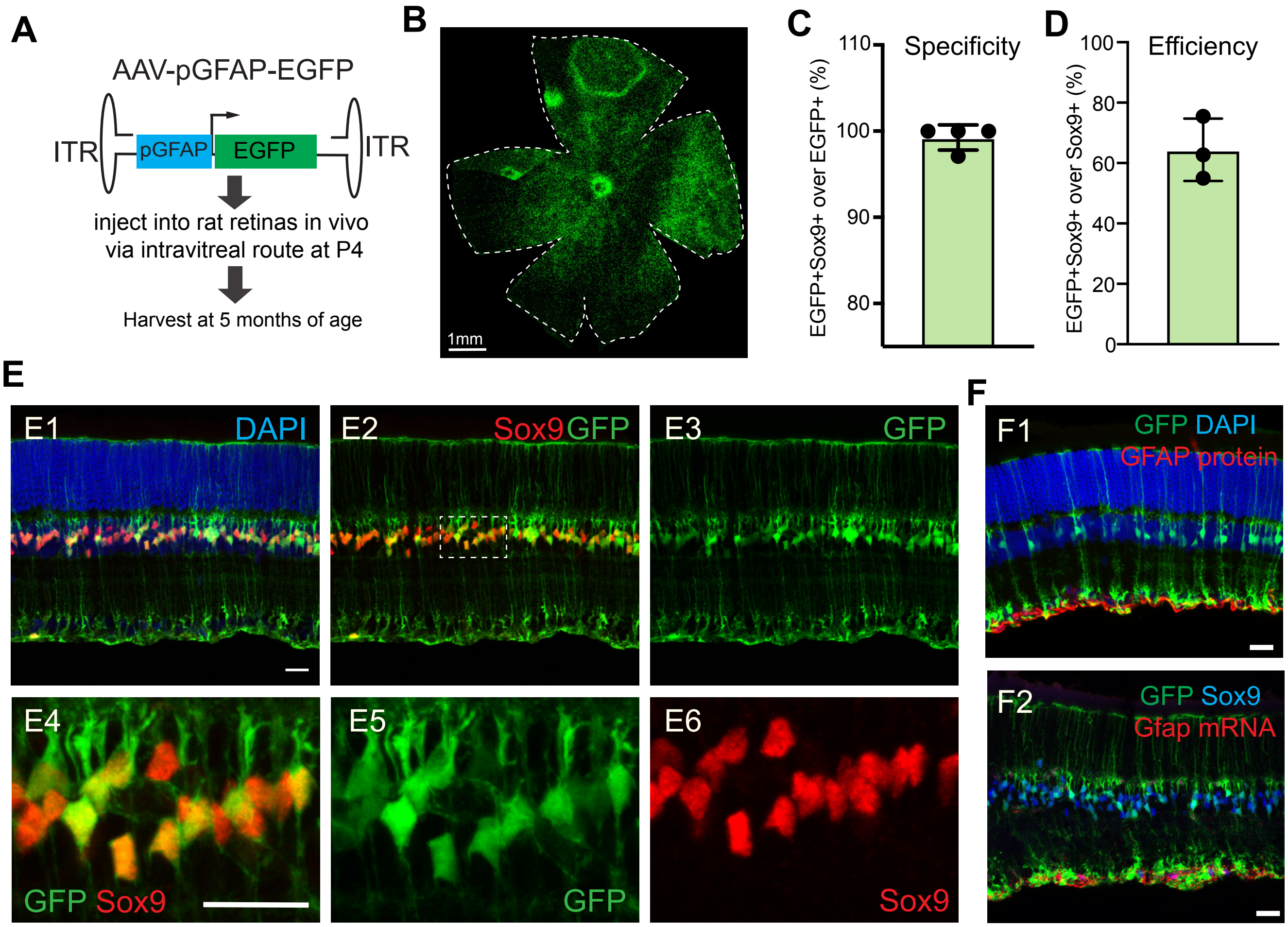


**Figure S7. The efficiency and specificity of AAV7m8-pGFAP-EGFP (related to Figure 4 and 5)**

**(A)** The AAV-pGFAP-EGFP construct and the experimental schematic design. **(B)** A representative image showing the transduction efficiency of AAV7m8-pGFAP-EGFP. Scale bar: 1mm. **(C-D)** The specificity (C) and efficiency (D) of AAV7m8-pGFAP-EGFP in labeling rat Müller glial cells. **(E)** Representative images showing the specificity and efficiency of AAV7m8-pGFAP-EGFP. Red: Sox9 antibody staining signals (Müller glial cell marker). Scale bar: 20um. **(F)** The *Gfap* mRNA and protein are not induced upon injection of AAV7m8-pGFAP-EGFP into the retina. Scale bar: 20um.

Figure S8

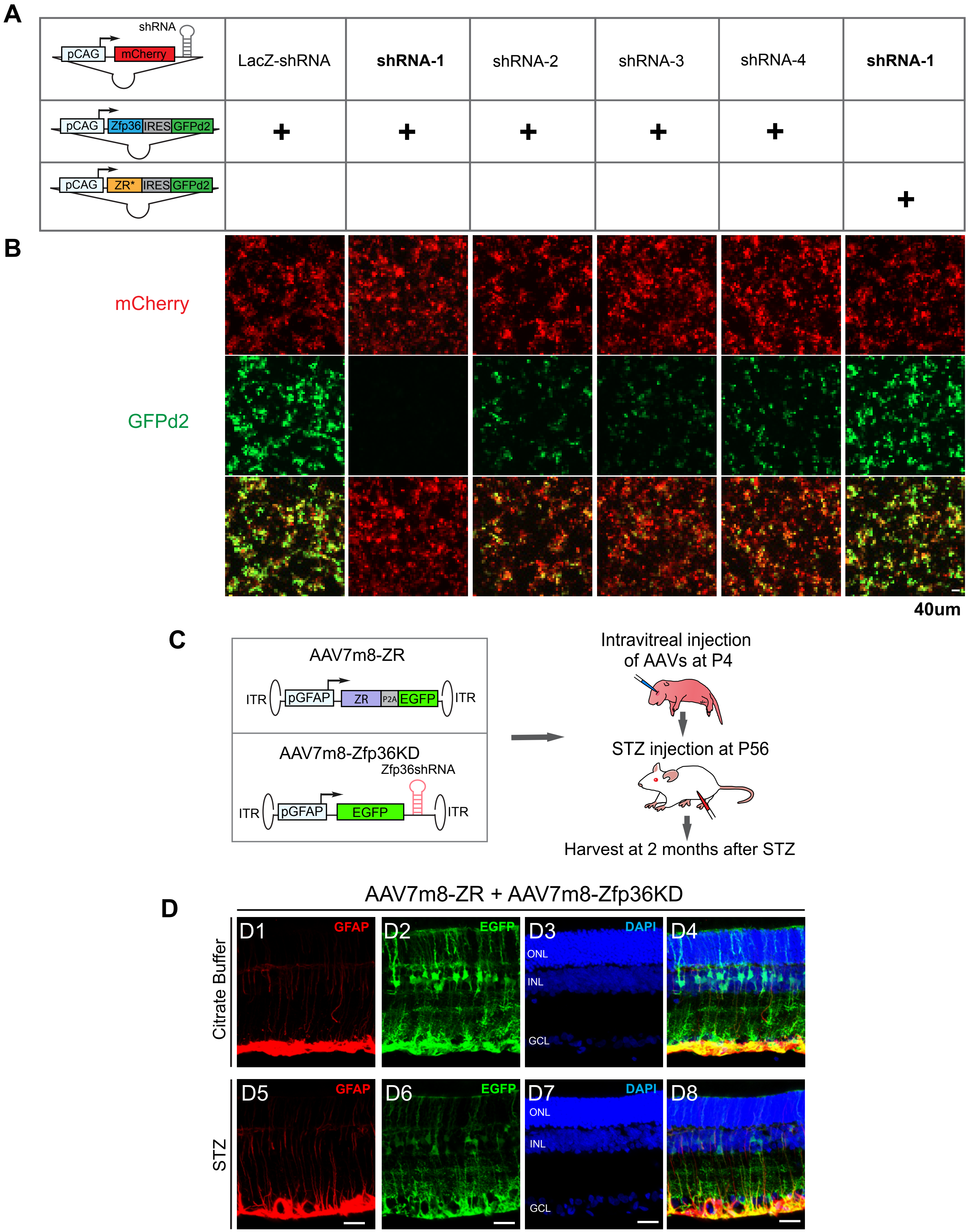
A
shRNA
pCAG
mCherry
LacZ-shRNA
shRNA-1
shRNA-2
shRNA-3
shRNA-4
shRNA-1
pCAG
Zfp36
IRES
GFPd2
+
+
+
+
+
pCAG
ZR*
IRES
GFPd2
+
B
mCherry
GFPd2
40um
C
AAV7m8-ZR
ITR
pGFAP
ZR
P2A
EGFP
ITR
AAV7m8-Zfp36KD
Zfp36shRNA
ITR
pGFAP
EGFP
ITR
Intravitreal injection
of AAVs at P4
STZ injection at P56
Harvest at 2 months after STZ
D
AAV7m8-ZR + AAV7m8-Zfp36KD
Citrate Buffer
D1
GFAP
D2
EGFP
D3
DAPI
ONL
INL
GCL
D4
STZ
D5
GFAP
D6
EGFP
D7
DAPI
ONL
INL
GCL
D8

**Figure S8. The knocking down efficiency of Zfp36-shRNA and its rescue by AAV7m8-ZR (related to Figure 4)**

**(A-B)** The experimental design schematic (A). shRNA-76 can efficiently knock down Zfp36 expression in HEK 293T cells in vitro (B). The ZR construct rescues this effect. GFPd2: destabilized form of GFP. **(C-D)** The experimental design schematic (C). The representative images showing retinas received AAV7m8-ZR and AAV7m8-Zfp36KD with citrate buffer or STZ injection (D). Red: GFAP staining. Scale bar: 20um.

**Table S1. The primers and DNA sequences used in the study**

| Name | Sequence (5’-3’) | Experiments |
|---|---|---|
| Zfp36-shRNA-1 | TGGAGGCTTGCTGAAGGCtgtaTGCTGATCGACATAAGGCTCTCGTAGGTTTTGGCCACT<br>GACTGACCTACGAGACTTATGTCGATCAGGACACAAGGCCTGTTACTAGCACTCACATG<br>GAACAAAT | shRNA with flanking sequences |
| ZR-p2A-EGFP | ATGGCCATTCGCGCCACCATGGATCTCTCTGCCATatatgaatctctcatgagcatGAGCCATGACCTGTCA<br>CCCGACCACGGAGGAACTGAGTCCTCCGGAGGACTTTGGAACATAAACTCATCGGACTCCATCC<br>CATCTGGGGTCACCTCTCGCCTGACTGGCCGCTCCACTAGCCTGGTGGAGGGCCGAAGCTGCA<br>GCTGGGTACCCCCACCCCCTGGTTTCGCACCCTTGGCTCCCCGCCCGGGCCCTGAGCTGTCAC<br>CCTCACCTACTTCGCCTACTGCAACTCCCACCACCTCCTCTCGATACAAGACTGAGCTCTGTCGG<br>ACCTACTCAGAGAGCGGGCGTTGTCGCTATGGGGCCAAGTGCCAGTTTGCCCACGGCCCGGGT<br>GAACTGCGCCAAGCCAATCGCCACCCCAAGTACAAAACGGAACTCTGCCACAAGTTCTACCTCCA<br>GGGCCGCTGCCCCTACGGCTCTCGATGCCACTTCATCCACAACCCTACCGAGGACCTGGCTCTC<br>CCTGGCCAGCCCCATGTGCTCCGACAAAGCATCAGCTTCTCAGGCTTGCCTTCAGGCCGCAGAA<br>CCTCACCACCACCTCCAGGCTTCTCTGGCCCTTCCCTGTCCTCTTGCTCCTTTTCGCCTTCCAGC<br>TCCCCACCACCGCCTGGGGACCTTCCACTTTCCCCTTCTGCCTTCTCTGCTGCCCCTGGGACCC<br>CTGTGTCTCGAAGAGACCCTACCCCAGCCTGTTGTCCCTCCTGCCGAAGGTCTACTACCCCTAG<br>CACCATCTGGGGGCCCTTGGGTGGTCTGGCTCGGAGCCCATCTGCACACTCTCTGGGATCCGA<br>CCCTGATGATTACGCCAGCAGCGGCAGCAGCCTGGGTGGGTCAGACTCGCCTGTCTTTGAGGC<br>CGGGGTGTTTGGGCCTCCTCAGCCCCCTGCACCCCCAAGGCGTCTTCCCATCTTCAATCGCATC<br>TCTGTCTCTGAGggaagcggagctactaacttcagcctgctgaagcaggctggagacgtggaggagaaccctggacctgctagcAT<br>GGTGAGCAAGGGCGAGGAGCTGTTCACCGGGGTGGTGCCCATCCTGGTCGAGCTGGACGGCG<br>ACGTAAACGGCCACAAGTTCAGCGTGTCCGGCGAGGGCGAGGGCGATGCCACCTACGGCAAGC<br>TGACCCTGAAGTTCATCTGCACCACCGGCAAGCTGCCCGTGCCCTGGCCCACCCTCGTGACCAC<br>CCTGACCTACGGCGTGCAGTGCTTCAGCCGCTACCCCGACCACATGAAGCAGCACGACTTCTTC<br>AAGTCCGCCATGCCCGAAGGCTACGTCCAGGAGCGCACCATCTTCTTCAAGGACGACGGCAACT<br>ACAAGACCCGCGCCGAGGTGAAGTTCGAGGGCGACACCCTGGTGAACCGCATCGAGCTGAAGG<br>GCATCGACTTCAAGGAGGACGGCAACATCCTGGGGCACAAGCTGGAGTACAACTACAACAGCCA<br>CAACGTCTATATCATGGCCGACAAGCAGAAGAACGGCATCAAGGTGAACTTCAAGATCCGCCAC<br>AACATCGAGGACGGCAGCGTGCAGCTCGCCGACCACTACCAGCAGAACACCCCCATCGGCGAC<br>GGCCCCGTGCTGCTGCCCGACAACCACTACCTGAGCACCCAGTCCGCCCTGAGCAAAGACCCC<br>AACGAGAAGCGCGATCACATGGTCCTGCTGGAGTTCGTGACCGCCGCCGGGATCACTCTCGGC<br>ATGGACGAGCTGTACAAGTAA | GSG_P2A_AS is highlighted by underlining. |
| Zfp36-p2A-EGFP | ATGGCCATTCGCGCCACCATGGATCTCTCTGCCATCTACGAGAGCCTTATGTCGATGAGCCATG<br>ACCTGTCACCCGACCACGGAGGAACTGAGTCCTCCGGAGGACTTTGGAACATAAACTCATCGGA<br>CTCCATCCCATCTGGGGTCACCTCTCGCCTGACTGGCCGCTCCACTAGCCTGGTGGAGGGCCG<br>AAGCTGCAGCTGGGTACCCCCACCCCCTGGTTTCGCACCCTTGGCTCCCCGCCCGGGCCCTGA<br>GCTGTCACCCTCACCTACTTCGCCTACTGCAACTCCCACCACCTCCTCTCGATACAAGACTGAGC<br>TCTGTCGGACCTACTCAGAGAGCGGGCGTTGTCGCTATGGGGCCAAGTGCCAGTTTGCCCACG<br>GCCCGGGTGAACTGCGCCAAGCCAATCGCCACCCCAAGTACAAAACGGAACTCTGCCACAAGTT<br>CTACCTCCAGGGCCGCTGCCCCTACGGCTCTCGATGCCACTTCATCCACAACCCTACCGAGGAC<br>CTGGCTCTCCCTGGCCAGCCCCATGTGCTCCGACAAAGCATCAGCTTCTCAGGCTTGCCTTCAG<br>GCCGCAGAACCTCACCACCACCTCCAGGCTTCTCTGGCCCTTCCCTGTCCTCTTGCTCCTTTTCG<br>CCTTCCAGCTCCCCACCACCGCCTGGGGACCTTCCACTTTCCCCTTCTGCCTTCTCTGCTGCCC<br>CTGGGACCCCTGTGTCTCGAAGAGACCCTACCCCAGCCTGTTGTCCCTCCTGCCGAAGGTCTAC<br>TACCCCTAGCACCATCTGGGGGCCCTTGGGTGGTCTGGCTCGGAGCCCATCTGCACACTCTCTG<br>GGATCCGACCCTGATGATTACGCCAGCAGCGGCAGCAGCCTGGGTGGGTCAGACTCGCCTGTC<br>TTTGAGGCCGGGGTGTTTGGGCCTCCTCAGCCCCCTGCACCCCCAAGGCGTCTTCCCATCTTCA<br>ATCGCATCTCTGTCTCTGAGggaagcggagctactaacttcagcctgctgaagcaggctggagacgtggaggagaaccctgga<br>cctgctagcATGGTGAGCAAGGGCGAGGAGCTGTTCACCGGGGTGGTGCCCATCCTGGTCGAGCTG<br>GACGGCGACGTAAACGGCCACAAGTTCAGCGTGTCCGGCGAGGGCGAGGGCGATGCCACCTAC<br>GGCAAGCTGACCCTGAAGTTCATCTGCACCACCGGCAAGCTGCCCGTGCCCTGGCCCACCCTC<br>GTGACCACCCTGACCTACGGCGTGCAGTGCTTCAGCCGCTACCCCGACCACATGAAGCAGCAC<br>GACTTCTTCAAGTCCGCCATGCCCGAAGGCTACGTCCAGGAGCGCACCATCTTCTTCAAGGACG<br>ACGGCAACTACAAGACCCGCGCCGAGGTGAAGTTCGAGGGCGACACCCTGGTGAACCGCATCG<br>AGCTGAAGGGCATCGACTTCAAGGAGGACGGCAACATCCTGGGGCACAAGCTGGAGTACAACTA<br>CAACAGCCACAACGTCTATATCATGGCCGACAAGCAGAAGAACGGCATCAAGGTGAACTTCAAG<br>ATCCGCCACAACATCGAGGACGGCAGCGTGCAGCTCGCCGACCACTACCAGCAGAACACCCCC<br>ATCGGCGACGGCCCCGTGCTGCTGCCCGACAACCACTACCTGAGCACCCAGTCCGCCCTGAGC<br>AAAGACCCCAACGAGAAGCGCGATCACATGGTCCTGCTGGAGTTCGTGACCGCCGCCGGGATC<br>ACTCTCGGCATGGACGAGCTGTACAAGTAA | GSG_P2A_AS is highlighted by underlining. |
